# The dithiocarbamate pesticides maneb and mancozeb disturb the metabolism of lipids and xenobiotics in an in vitro model of metabolic dysfunction-associated steatotic liver disease

**DOI:** 10.1101/2024.05.16.594496

**Authors:** Kilian Petitjean, Giovanna Dicara, Sébastien Bristeau, Hugo Coppens-Exandier, Laurence Amalric, Nicole Baran, Camille C. Savary, Anne Corlu, Pascal Loyer, Bernard Fromenty

**Author notes:** Corresponding author: Bernard Fromenty. **e-mail addresses of all authors:** Kilian Petitjean, Giovanna Dicara, Sébastien Bristeau, Hugo Coppens-Exandier Laurence Amalric, Nicole Baran, Camille Savary Anne Corlu, Pascal Loyer, Bernard Fromenty.

## Abstract

Pesticides are increasingly recognized to be hepatotoxic but less is known about their toxicity in metabolic dysfunction-associated steatotic liver disease (MASLD). Recent investigations reported oxidative stress-induced apoptosis in differentiated hepatocyte-like HepaRG cells after a single treatment with a 7-pesticide mixture that included chlorpyrifos, dimethoate, diazinon, iprodione, imazalil, and the dithiocarbamates maneb and mancozeb. These effects were reproduced by maneb, mancozeb, or manganese chloride (MnCl_2_). Herein, differentiated HepaRG cells cultured for 2 weeks without (-FA) or with (+FA) a mixture of stearic and oleic acids were treated with this 7-pesticide mixture, maneb, mancozeb, or MnCl_2_ along the same period. While these molecules did not induce neutral lipid accumulation in -FA-HepaRG cells, they worsened steatosis in +FA-HepaRG cells. Maneb or MnCl_2_ impaired very low-density lipoprotein (VLDL) secretion and increased fatty acid uptake without altering mitochondrial fatty acid oxidation and *de novo* lipogenesis. Reduced VLDL secretion was associated with decreased mRNA levels of apolipoproteins B and C3 and microsomal triglyceride transfer protein. Zinc supplementation restored VLDL secretion, reduced fatty acid uptake and prevented the exacerbation of steatosis in +FA-HepaRG cells treated with mancozeb or MnCl_2_. The mixture, maneb, or MnCl_2_ also reduced the mRNA expression and activity of several cytochromes P450 in +FA- and -FA-HepaRG cells. This was associated with impaired biotransformation of diazinon while chlorpyrifos metabolism was unaffected. Hence, maneb, mancozeb and MnCl_2_ disturb the metabolism of lipids and xenobiotics in HepaRG cells, in particular in fatty acid-exposed cells. These findings could have major pathophysiological consequences in dithiocarbamate-exposed individuals with MASLD.

## Introduction

Occupational or environmental exposure to different pesticides is suspected to increase the risk of developing several types of metabolic disturbances including overweight and obesity (Araújo et al. 2022; Zuo et al. 2022), metabolic syndrome (Lamat et al. 2022), type 2 diabetes (Evangelou et al. 2016; Rebouillat et al. 2022) and hepatic steatosis (Sang et al. 2022; Wahlang et al. 2020), also referred to as fatty liver. Of note, pesticide exposure can induce liver lesions other than steatosis such as hepatic cytolysis, cholestasis and different types of vascular disorders (EASL 2019; Vilas-Boas et al. 2019).

Notably, steatosis is very frequent in obese individuals (Brunt et al. 2021) and in alcoholic patients (Arab et al. 2023). It can also occur in a significant number of cases of drug-induced liver injury (Fromenty 2019). A recent nomenclature introduced the term steatotic liver disease (SLD) which encompasses all types of fatty liver disease regardless of their cause (Rinella et al. 2023). In this nomenclature, steatotic liver disease in obese individuals is referred to as metabolic dysfunction-associated steatotic liver disease (MASLD) (Rinella et al. 2023).

MASLD in obese individuals can be worsened by alcohol and some xenobiotics (Fromenty and Roden 2023; Klaunig et al. 2018; Massart et al. 2022). Hence, many endogenous and exogenous factors can interact with each other to favor the occurrence or the worsening of steatotic liver disease (Klaunig et al. 2018; Larrain and Rinella 2012; Massart et al. 2022). These multiple interactions are worrisome since many overweight or obese people are treated with different pharmaceuticals and/or exposed to a variety of environmental pollutants. Even more worrying could be the progression of steatosis to advanced chronic liver disease in a significant number of exposed individuals (Tovoli et al. 2023). Indeed, although steatosis is a benign condition in the short term regardless of its cause, it can progress in the long term into steatohepatitis, advanced fibrosis, cirrhosis and hepatocellular carcinoma (Arab et al. 2023; Brunt et al. 2021; Fromenty 2019).

There is now ample evidence that obesity and MASLD are associated with altered hepatic activity of different xenobiotic-metabolizing enzymes (XMEs) such as cytochromes P450 (CYPs) and uridine diphosphate glucuronosyltransferases (UGTs) (Brill et al. 2012; Cobina and Akhlagi 2017; Smit et al. 2018). In particular, increased CYP2E1 activity and reduced CYP3A4 activity have been consistently reported in obesity and MASLD (Begriche et al. 2023; Brill et al. 2012; Bucher et al. 2018; Cobina and Akhlagi 2017; Smit et al. 2018). The mechanisms of XME alterations in these metabolic diseases are still unclear and seem to be complex. For instance, increased CYP2E1 activity might be linked to hyperleptinemia, hyperglucagonemia and insulin resistance and the intake of some saturated fatty acids (Aubert et al. 2011; Begriche et al. 2023; Massart et al. 2021). Nevertheless, there is currently no information about the precise downstream signaling pathways involved in CYP2E1 induction during obesity and MASLD. Interestingly, MASLD also seems to be associated with altered hepatic activity of several nuclear receptors involved in xenobiotic metabolism such as the aryl hydrocarbon receptor (AhR), constitutive androstane receptor (CAR) and pregnane X receptor (PXR) (Cave et al. 2016; Klaunig et al. 2018; Yang et al. 2020). Notably, these nuclear receptors are also involved in lipid and carbohydrate metabolism and recent investigations unveiled their possible role in MASLD pathogenesis (Cave et al. 2016; Klaunig et al. 2018; Yang et al. 2020). Again, these data demonstrate a close relationship between xenobiotic biotransformation and intermediary metabolism in the liver.

In a recent study carried out in metabolically competent differentiated HepaRG cells and other hepatic cells, we investigated the acute cytotoxic effects of 7 pesticides, namely chlorpyrifos-ethyl (henceforth referred to as chlorpyrifos), dimethoate, diazinon, iprodione, imazalil, maneb and mancozeb (Petitjean et al. 2024), which are often detected in food samples in many countries (Caldas et al. 2004; EFSA 2017; Ssemugabo et al. 2022). Notably, our data showed that the manganese (Mn)-containing dithiocarbamates (DTCs) maneb and mancozeb were solely responsible for the acute cytotoxicity induced by the mixture (Petitjean et al. 2024). Moreover, our investigations unveiled that maneb and mancozeb-induced reactive oxygen species (ROS) overproduction and apoptosis could be attributed to the release of Mn leading to intracellular Mn overload and depletion in zinc (Zn). Lastly, maneb and mancozeb-induced cytotoxicity was prevented by Zn supplementation, thus showing that Mn-containing fungicides are toxic by disrupting Mn and Zn homeostasis, thus triggering oxidative stress and cell death in human hepatocytes (Petitjean et al. 2024).

In this study, we wished to proceed with our investigations to determine whether repeated exposure to this 7-pesticide mixture could impair lipid and xenobiotic metabolism in differentiated HepaRG cells. Because MASLD can be worsened by some xenobiotics (Fromenty and Roden 2023; Klaunig et al. 2018; Massart et al. 2022), we investigated the effects of these pesticides in HepaRG cells cultured or not with a mixture of stearic and oleic acids. Importantly, we previously showed that culturing HepaRG cells with these fatty acids brought about a valuable *in vitro* model of MASLD, which reproduced different metabolic features of the human liver disease including higher CYP2E1 and reduced CYP3A4 activities (Bucher et al. 2018; Le Guillou et al. 2018). We also used this cellular model to investigate hydroxychloroquine biotransformation (Ferron et al. 2021) and SARS-CoV-2 receptor ACE2 expression (Cano et al. 2023) in the setting of MASLD.

## Materials and Methods

### Chemicals and reagents

Chlorpyrifos, dimethoate, diazinon, imazalil, iprodione, maneb, mancozeb, dimethyl sulfoxide (DMSO), oleic acid, stearic acid, palmitic acid, palmitoyl-CoA, palmitoyl-L-carnitine, fatty acid-free bovine serum albumin (BSA), manganese chloride (MnCl_2_) and zinc chloride (ZnCl_2_) were purchased from Sigma-Aldrich (St. Louis, MO, USA). The Chemical Abstracts Service (CAS) numbers of the 7 pesticides are provided in Table 1. William’s E medium, Dulbecco’s phosphate-buffered saline (PBS) Gibco™, glutamine, penicillin, streptomycin, formaldehyde, Nile red and Hoechst 33342 dyes were obtained from Thermo Fischer Scientific (Waltham, MA). Radiolabeled [^14^C(U)]palmitic acid (31.45 GBq/mmol, 3.7 MBq/mL) and [2-^14^C] acetic acid (2.183 GBq/mmol, 37 MBq/mL) were purchased from PerkinElmer (Waltham, MA). For the ultra[performance liquid chromatography-tandem mass spectrometry (UPLC-MS/MS) analyses, analytical internal standards of diazinon (deuterated diazinon D_10,_ 99.7%), 2-isopropyl-6-methyl-4-pyrimidinol (IMP ^13^C_4_, 99.9%), chlorpyrifos (deuterated chlorpyrifos D_10_, 99.9%) and 3,5,6-trichloro-2-pyridinol (TCP ^13^C_3_, 99.3%) were purchased from Techlab (Saint-Julien-lès-Metz, France). Hydrocortisone hemisuccinate was from Pharmacia & Upjohn (Guyancourt, France). DMSO was used as a solvent for the 7 pesticides to prepare a stock solution of 100 mM, except for diazinon for which the concentration was 10 mM. In this study, the term “pesticide mixture” is referred to as the combination of the 7 compounds.

**Table 1.**
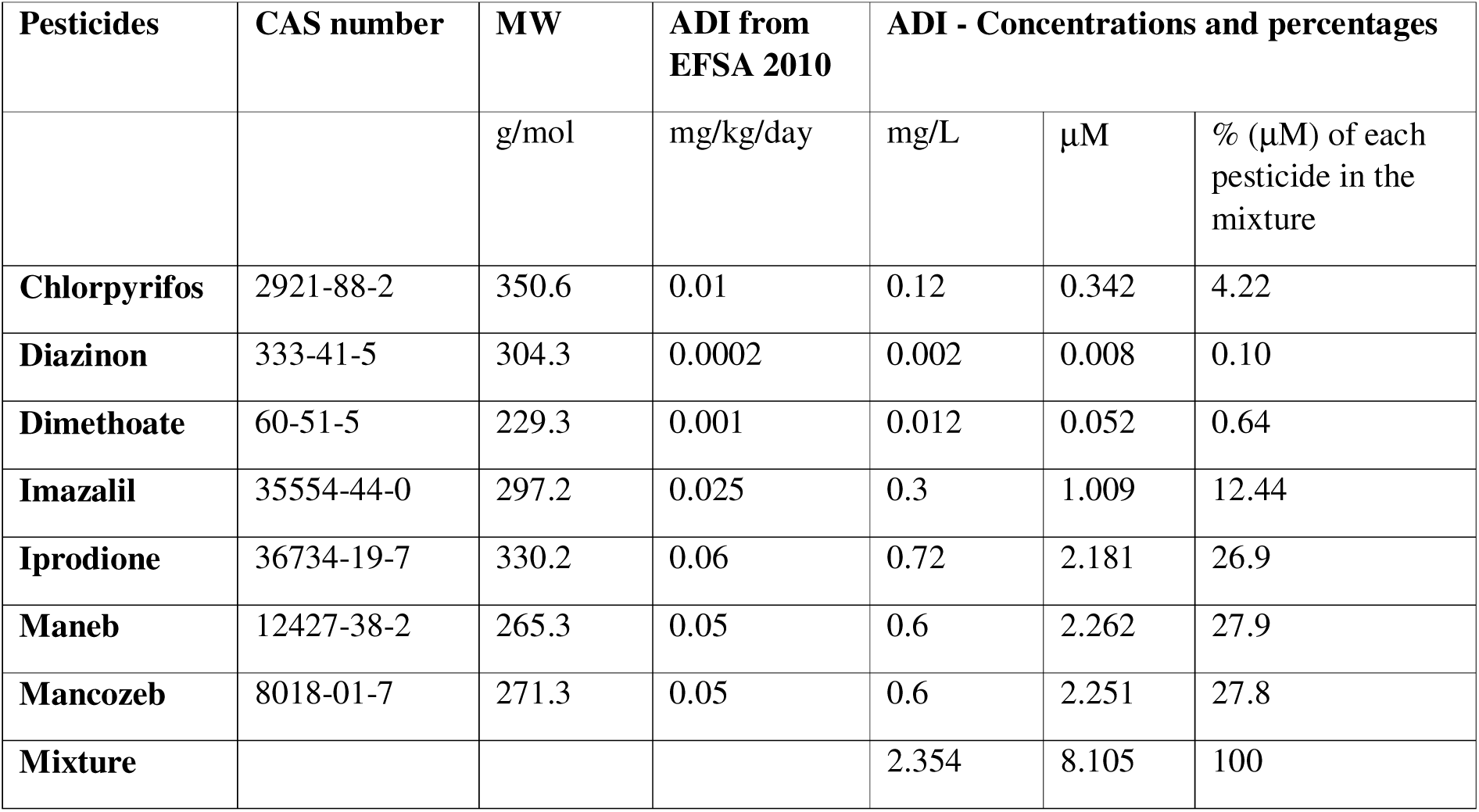
CAS number, molecular weight (MW) and ADI of the different pesticides and the mixture.

### Cell culture and treatments

Native HepaRG cells were cultured as previously described, with minor modifications (Cerec et al. 2007). First, progenitor HepaRG cells were seeded at 2.6x10^4^ cells/cm^2^ and were incubated in William’s E Medium supplemented with 10% fetal bovine serum (FBS) (Biosera^TM^ S006R20001, Hyclone^TM^ SH30066.03, *v/v*:3/1), 1% glutamine (200 mM), 100 U/ml penicillin, 100 µg/ml streptomycin, 5 µg/ml insulin, and 50 µM hydrocortisone hemisuccinate. After 2 weeks, cell differentiation was enhanced with 2% DMSO added to the medium for 2 additional weeks. Two days before treatments, FBS and DMSO concentrations were lowered to 5% and 1%, respectively as previously described (Bucher et al. 2018). Then, differentiated HepaRG cells were exposed every 2 or 3 days for 2 weeks to a medium containing or not stearic and oleic acids (150 µM each) and the pesticides. HepaRG cells cultured or not with stearic and oleic acids will hereafter be referred to as +FA-HepaRG and -FA-HepaRG cells, respectively. All investigations were performed 24 hours (h) after the last treatment. Control conditions corresponded to HepaRG cells treated with 1% DMSO without fatty acids and pesticides. HepaRG cells were always maintained in incubators at 37 °C with 5% CO_2_ and saturating humidity. Investigations were carried out on passages 12 to 17. The concentrations of pesticides used to treat the cells were extrapolated from the international Acceptable Daily Intake (ADI) (EFSA 2010), which represents the threshold of safety in humans for lifetime exposure (Hochane et al. 2017; Petitjean et al. 2024). Briefly, the ADI value (mg/kg.bw/day) for each pesticide was converted to mg/L considering the total absorption of the ingested amount and then its dilution in 5 liters of blood (Hochane et al. 2017; Petitjean et al. 2024). The concentration for each pesticide was then converted to μM (Table 1).

In a few investigations, two different batches of primary human hepatocytes (PHH) were used to confirm some results obtained in HepaRG cells. These batches were obtained from the Centre de Ressources Biologiques (CRB) Santé of Rennes (institutional review board approval BB-0033-00056). Information regarding the donors for PHH 1 and PHH 2 is provided in the Supplemental material (Table 1). Briefly, PHH were seeded at a density of 3x10^5^ cells/cm^2^ in the same medium as progenitor HepaRG cells. When cells were adherent, the medium was replaced by the same medium as differentiated HepaRG cells. One day before treatments, FBS and DMSO concentrations were lowered to 5% and 1%, respectively. PHH were exposed to fatty acids and treated with maneb, mancozeb, or MnCl_2_ using the same protocol as for HepaRG cells, except for time exposure which was shortened to 10 or 7 days for PHH 1 and PHH 2, respectively. Indeed, PHH cultures had to be stopped before significant cell detachment, which was heralded by subtle changes in PHH morphology.

### Measurement of cellular ATP levels

Cell viability was assessed by measuring the relative intracellular ATP levels using the CellTiter-Glo Luminescent Cell viability assay (Promega, Charbonnières-les-Bains, France). After treatment with pesticides, cells were incubated with the CellTiter-Glo reagent for 10 min. Lysates were transferred to an opaque multi-well plate and luminescent signals were quantified at 540 nm using a POLARstar Omega microplate reader (BMG Labtech, Ortenberg, Germany). Cell viability in treated cells was expressed as the percentage of the luminescent values measured in untreated cells, which was arbitrarily set as 100%.

### Assessment of neutral lipids and triglyceride quantification

Accumulation of neutral lipids was evaluated using the Nile red dye (Greenspan et al. 1985), as recently described with minor modifications (Bucher et al. 2018). Briefly, HepaRG cells were washed 3 times with PBS, fixed and stained with PBS containing 4% formaldehyde and 10 µg/mL Hoechst 33342 dye for 30 min. After 3 washes with PBS, cells were incubated in PBS containing 0.1 µg/mL Nile red for 30 min. Then, cells were washed 3 times with PBS and fluorescence intensities were measured using a POLARstar Omega microplate reader (BMG Labtech) with excitation/emission wavelengths at 355 /460 nm and 520/590 nm for Hoechst 33342 and Nile red, respectively. Nile red fluorescence intensity values were normalized to Hoechst 33342 fluorescence intensity. Cellular triglyceride levels were quantified using the Triglyceride-Glo^TM^ assay (Promega) and then normalized to the concentration of total proteins assessed by using the Pierce™ BCA Protein Assay Kit (Thermo Fisher Scientific).

### Assessment of fatty acid uptake

Fatty acid uptake was measured by using the Fatty Acid Uptake kit from Sigma-Aldrich. Briefly, HepaRG cells were washed with warm PBS and incubated at 37°C for 1 h in an unsupplemented William’s E medium. Cells were then washed with warm PBS and incubated with a fluorescent probe derived from dodecanoic acid (TF2-C12) for 1.5 h at 37°C. After 2 washes with PBS, fluorescence intensity was measured using a POLARstar Omega microplate reader (BMG Labtech) with excitation/emission wavelengths at 482/520 nm. TF2-C12 fluorescence intensity values were normalized to total protein content and expressed in comparison to control cells.

### Assessment of *de novo* lipogenesis

*De novo* lipogenesis (DNL) in HepaRG cells was assessed by measuring newly synthesized radiolabeled lipids from [2-^14^C] acetic acid, as previously described (Allard et al. 2021). Briefly, the culture medium was removed and cells were washed with warm PBS. Next, cells were incubated for 3 h with phenol red-free William’s E medium containing 1% fatty acid-free BSA, 50 µM cold acetic acid and [2-^14^C]acetic acid (1.85 kBq per well, corresponding to 16.9 nM). The medium was then removed and cells were washed with warm PBS before adding a mixture of hexane/isopropanol (3/2;v/v). Cell culture plates were sealed and incubated for 1 h at room temperature for lipid extraction. After transfer of the plate content in 200 μL microtubes, hexane and PBS were added to have a hexane/isopropanol/PBS ratio of 6/2/3 (v/v/v). Microtubes were then centrifuged at 1000*g* for 5 min and radiolabeled lipids were counted in the upper phase with a Tri-Carb 4910TR liquid scintillation counter (PerkinElmer). Results were normalized to total protein content and expressed in comparison to control cells.

### Assessment of mitochondrial fatty acid oxidation

Mitochondrial fatty acid oxidation (FAO) was assessed by measuring the acid-soluble radiolabeled metabolites resulting from the mitochondrial oxidation of [^14^C(U)]palmitic acid, as recently described (Allard et al. 2021). In brief, the culture medium was removed and cells were washed with warm PBS before the addition of phenol red-free William’s E medium containing 1% fatty acid-free BSA, 100 µM cold palmitic acid, [^14^C(U)]palmitic acid (185 Bq per well, corresponding to 117 pM), 1 mM L-carnitine and 1% DMSO. After 3 h of incubation at 37 °C, perchloric acid (final concentration, 6%) was added and plates were centrifuged at 2,000g for 10 min, and the supernatant was counted for [^14^C]-labeled acid-soluble β-oxidation products using a Tri-Carb 4910TR liquid scintillation counter (PerkinElmer). These [^14^C]-labeled acid-soluble β-oxidation products are deemed to be mainly ketone bodies and to a lesser extent intermediates of the tricarboxylic acid cycle (Fromenty et al. 1993). Results were normalized to total protein content and expressed in comparison to control cells.

### Assessment of very low-density lipoprotein secretion

Secretion of very low-density lipoprotein (VLDL) from hepatic cells can be assessed via the measurement of apolipoprotein B 100 (hereafter referred to as apoB) in cell supernatant (Allard et al. 2021; Grünig et al. 2018). Hence, apoB levels were measured in HepaRG cell supernatants using the Human Apolipoprotein B ELISA Pro kit from Mabtech (Nacka Strand, Sweden) according to the manufacturer’s instructions, as recently described (Allard et al. 2021). Briefly, cell supernatants were centrifuged at 1500*g* for 5 min and diluted to ¼ in Apo ELISA buffer. Samples were then incubated in pre-coated strip plates provided in the kit. After 2 h incubation, samples were removed and plates were washed and incubated for 1 h with monoclonal anti-human apoB (LDL-11-biotin) antibody. Plates were then washed and incubated with a streptavidin-horseradish peroxidase-conjugated antibody. After the last washing, 3,3’,5,5’-tetramethylbenzidine (TMB) was used as a substrate for 15 min, protected from light. The reaction was stopped and absorbance was measured at 450 nm using a POLARstar Omega microplate reader (BMG Labtech). Finally, apoB levels were calculated utilizing a 4-parameter standard curve and expressed in comparison to control cells.

### Measurement of CYP, UGT and sulfotransferase activities

CYP2E1 activity was measured by determining the generation of chlorzoxazone O-glucuronide from chlorzoxazone with a high-performance liquid chromatography (HPLC) method using an Accucore PFP column (Thermo Scientific, Waltham, MA) (Bucher et al. 2018; Quesnot et al. 2018). Briefly, HepaRG cells were incubated for 6 h at 37°C in phenol red-free and DMSO-free William’s E medium containing 300 µM chlorzoxazone. At the end of the incubation, aliquots of culture medium were collected and chlorzoxazone O-glucuronide was measured by HPLC analysis with UV detection at 287 nm. CYP2E1 activity was expressed as pmol/mg protein/h.

CYP3A4/5, CYP2B6, CYP1A2, CYP2C19, UGT and sulfotransferase (SULT) activities were respectively measured by the production of hydroxymidazolam, hydroxybupropion, acetaminophen, 4’-hydroxymephenytoin, acetaminophen glucuronide and acetaminophen sulfate using liquid chromatography (LC) coupled to mass spectrometry (MS), as previously described (Anthérieu et al. 2010), with some modifications. Briefly, HepaRG cells were washed at the end of the 2-week treatments with warm PBS and incubated for 30 min in a red phenol-free William’s E medium at 37°C. This step aimed at removing all pesticide metabolites synthesized during the 2-week treatment. Next, the medium was replaced by a red phenol-free William’s E containing 100 µM midazolam, 100 µM bupropion, 200 µM phenacetin, 100 µM S-mephenytoin or 1 mM acetaminophen. After 1 h at 37°C, supernatants were then collected and diluted to ½ in acetonitrile in a 96-well plate and kept at -20°C until analysis. After 10 min shaking, samples were directly injected into the Exion LC AC system (SCIEX, Toronto, Canada). Compounds were separated on an Xselect HSS T3 column (2.1x75 mm, 2.5 µM, Waters, Milford, MA) maintained at 35°C. The mobile phase was composed of 2 mM ammonium formate containing 0.1% formic acid and acetonitrile. The total LC run was 5.7 min with a flow rate of 0.5 mL/min. Samples were kept at 4°C until injection. Detection was operated on the triple quadrupole mass spectrometer AB SCIEX Triple Quad 5500+ (SCIEX) equipped with an electrospray ion source in positive mode. Data processing was performed using Analyst 1.7.2 software (SCIEX). Activities were expressed as nmol/mg protein/h.

### Measurement of diazinon, IMP, chlorpyrifos and TCP concentrations

To determine whether HepaRG cell exposure to pesticides could disturb the biotransformation of diazinon to IMP and that of chlorpyrifos to TCP, levels of these 4 compounds were measured in the culture medium by ultra-performance LC-MS/MS (UPLC-MS/MS). Hence, at the end of the 2-week treatments with the pesticides, HepaRG cells were washed with warm PBS and incubated for 30 min in a red phenol-free William’s E medium at 37°C. This step aimed at removing all pesticide metabolites synthesized during the 2-week treatment. Next, the medium was replaced by a red phenol-free William’s E containing 50 µM of diazinon, or 50 μM of chlorpyrifos. After 8 and 24 h of incubation at 37°C, the cell culture medium was collected and HepaRG cells were washed with an equal volume of ultrapure water which was also collected. The resulting mixtures were centrifuged at 20,000*g* for 20 min at 4°C and the supernatants were then transferred and frozen until analysis. All compounds were simultaneously quantified in supernatant by UPLC-MS/MS with a Xevo-TQD ® triple quadrupole (Waters, Maryville, TN) in MRM (Multiple Reaction Monitoring) mode and with positive electrospray ionization. Chromatographic separation was done with a Waters Acquity UPLC HSS-T3 column (2.1 mm × 100 mm, particle size 1.8 µm) maintained at 30°C. The mobile phase was a gradient of water/0.01% formic acid v/v (A) and acetonitrile/0.01% formic acid v/v (B) with a flow rate of 0.3 mL/min. The gradient started with 95% eluent A, maintaining isocratic conditions for 0.5 minutes. Then eluent B increased to 100% in 6 minutes and was maintained for 0.50 minutes. Finally, initial conditions were reached again in 0.10 minutes with a re-equilibrium time of 2.9 minutes to restore the column. The injection volume was 2 µL. For the mass detector, the parameters were as follows: source temperature 150 °C, desolvation temperature 650 °C, cone gas flow 50 L/h, desolvation gas flow 800 L/h, positive ion mode (spray voltage 2.5 kV).

The injection of analytical standards (at a concentration of 10 µM in water) allowed us to define the analytical features of the compounds and to estimate their limit of quantification. Quantification was based on peak area and was performed using the 4 internal standards, namely deuterated diazinon D_10_, IMP ^13^C_4_, deuterated chlorpyrifos D_10_ and TCP ^13^C_3_. To determine the calibration function, injection of 9 concentrations (in a mixture of water/ William’s E medium (50/50 v/v)) was performed, ranging from 0.003 to 3 µM for diazinon and IMP and from 0.03 to 30 µM for chlorpyrifos and TCP. Data was best fitted with a quadratic calibration except for IMP (linear function). The limit of quantification was 0.03 µM for each compound. For each series, a control (0.05 µM for diazinon and its metabolite and 5 µM for chlorpyrifos and its metabolite) was injected every 30 samples. The concentrations of the different compounds were normalized by the total protein content. Of note, despite our efforts to measure chlorpyrifos concentrations in the collected biological samples, the quantification of this pesticide was not possible since its concentration was below the limit of quantification.

### mRNA expression studies

RNA purification and reverse transcription were performed using the Nucleospin RNA kit from Macherey-Nagel (Düren, Germany) and the High capacity cDNA reverse transcription kit from Applied Biosystems (Woolston, UK), respectively. Quantitative PCR was carried out with the Sybr Green PCR Master Mix from Applied Biosystems on an ABI PRISM 7900HT instrument. The primer sequences are provided in the Supplemental material (Table 2). The expression of TATA-box binding protein (TBP) was chosen as a reference and the 2^-ΔΔCt^ calculation method was used to express the relative expression of each selected gene.

### Statistical analysis

All data were expressed as mean ± standard error of the mean (SEM) from at least three independent experiments. GraphPad Prism 8 software (GraphPad Software Inc, San Diego, CA) was used for all statistical analyses. Most investigations were carried out by varying two factors, namely fatty acid exposure and treatments with different compounds. For each data set, the Shapiro-Wilk normality test was performed. When this test was positive, comparisons between multiple groups were performed using a two-way analysis of variance (ANOVA) followed by a *post hoc* Dunnett’s test or Sidak’s test, as recommended by the GraphPad software. For a few analyses with only one factor (i.e. pesticide treatment), a one-way ANOVA was used followed by a *post hoc* Dunnett’s test. When the data were not normally distributed, a Kruskal-Wallis test was performed followed by a *post hoc* Dunn’s test. The threshold for statistical significance was set to p < 0.05.

## Results

### Pesticide-induced cytotoxicity in HepaRG cells after a 2-week exposure

In our recent investigations, pesticide-induced cytotoxicity was assessed in hepatocyte-like HepaRG cells cultured with 2% DMSO (Petitjean et al. 2024). In the present study, DMSO concentration was lowered to 1% to curb this cytotoxicity since the primary objective of our investigations was to study lipid and xenobiotic metabolism in differentiated HepaRG cells in a culture condition of reduced cell death. Indeed, previous studies in HepaRG cells consistently showed that high DMSO concentrations close to 2% significantly enhance the expression and activity of different XMEs including CYPs (Aninat et al. 2006; Dubois-Pot-Schneider et al. 2022; Klein et al. 2014), which explained the previously reported higher pesticide-induced cytotoxicity in HepaRG cells (Petitjean et al. 2024).

In a preliminary series of experiments in -FA and +FA-HepaRG cells, the cytotoxicity of each pesticide or their combination was assessed after 2 weeks of exposure over a wide range of concentrations. Our results showed that at the highest studied concentration (i.e. 3ADI), the reduction of cellular ATP content was never above 30% in both -FA and +FA-HepaRG cells (Supplemental Fig. 1). This 30% cytotoxicity was brought about only by maneb and mancozeb while the other pesticides induced less or no cytotoxicity at the tested concentrations (Supplemental Fig. 1). Notably, the 7-pesticide mixture including maneb and mancozeb induced a concentration-dependent loss of ATP in - FA and +FA-HepaRG cells, while a mixture without these dithiocarbamates did not induce significant cytotoxicity (Supplemental Fig. 1). In addition, the combination of maneb + mancozeb (dithiocarbamates) induced cytotoxicity that was about similar to the one observed with each dithiocarbamate, or the mixture (Supplemental Fig. 1). Finally, MnCl_2_ cytotoxicity was also evaluated because this metal is released in differentiated HepaRG cells during maneb and mancozeb metabolism (Petitjean et al. 2024). However, no significant loss of ATP content was detected for 6.75 μM MgCl_2_ (Supplemental Fig. 1), which corresponds to the 3ADI concentration of maneb, or mancozeb (Table 1). Interestingly, these cytotoxicity data on repeated pesticide treatments (Supplemental Fig. 1) are in line with our previous study showing that among the 7 pesticides, only maneb and mancozeb brought about significant loss of ATP after acute exposure (Petitjean et al. 2024).

### Pesticide-induced accumulation of neutral lipids and triglycerides in differentiated HepaRG cells

Next, we wished to determine the steatogenic potential of the 7 pesticides in HepaRG cells using the Nile red dye (Fig. 1a), which allows for assessing the cellular content of neutral lipids (Allard et al. 2021; Bucher et al. 2018). For this purpose, HepaRG cells cultured for 2 weeks with or without a mixture of stearic and oleic acids were exposed or not to the pesticides along the same period. Five different concentrations of pesticides were studied, namely 1/15, 1/10, 1/5, 1 and 3ADI.

**Figure 1.**
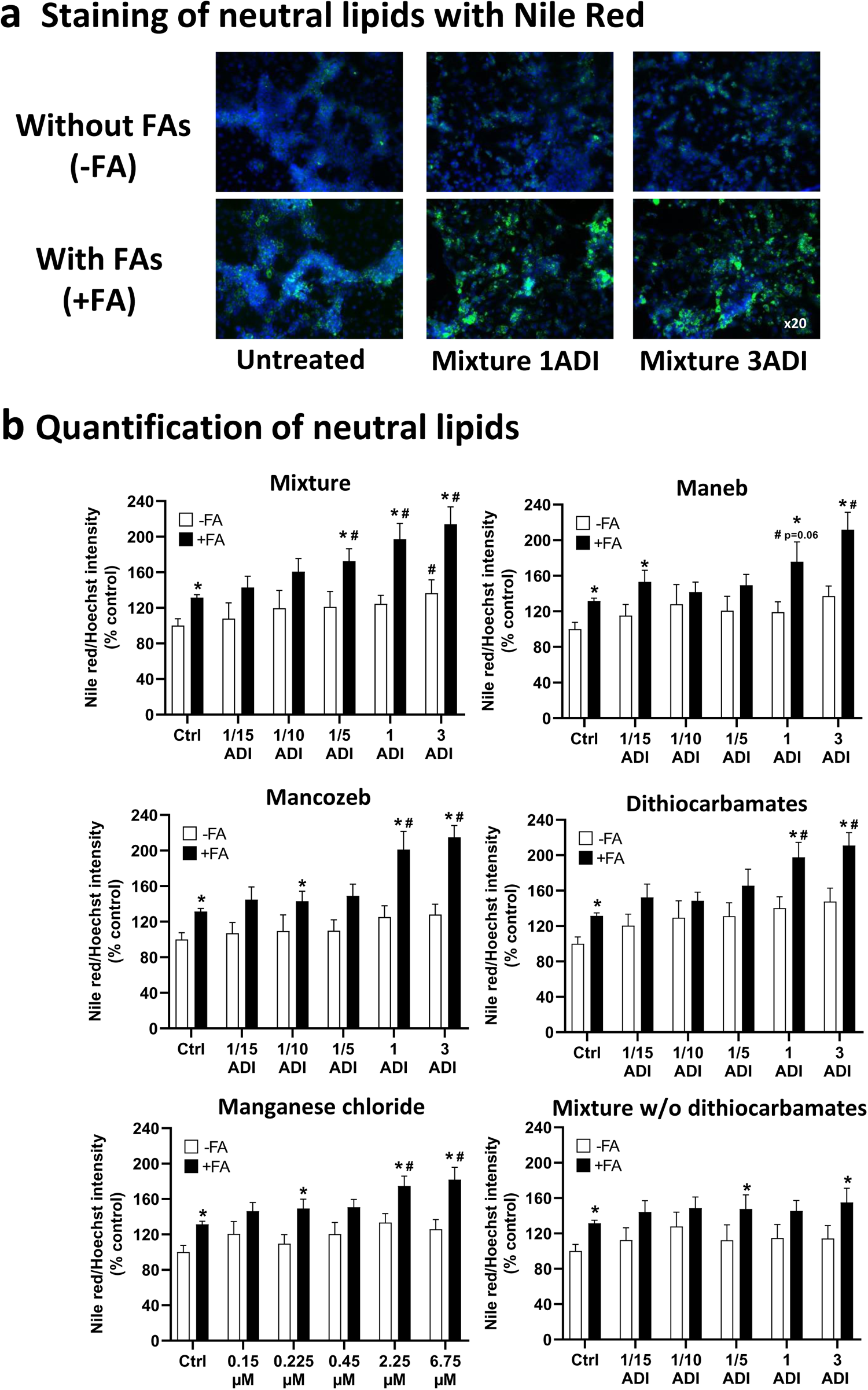
Effects of pesticides and manganese chloride on neutral lipid accumulation in differentiated HepaRG cells cultured with or without stearic and oleic acids. HepaRG cells were treated with (+FA) or without (-FA) fatty acids, pesticides, or manganese chloride (MnCl_2_) for 2 weeks. Pesticide concentrations ranged from 1/15 to 3 ADI, while those of MnCl_2_ ranged from 0.15 to 6.75 μM, corresponding to the respective ADI concentrations of maneb and mancozeb. Neutral lipids and nuclei were stained with the fluorescent dyes Nile red and Hoechst 33342, respectively. (**a**) Representative pictures of -FA and +FA-HepaRG cells exposed or not to the pesticide mixture at the ADI and 3ADI concentrations. Neutral lipids and nuclei are visualized in green and blue, respectively. Pictures were taken at a magnification of ×20. (**b**) Quantification of neutral lipids in -FA and +FA-HepaRG cells treated with pesticides or MnCl_2_ using the Nile red/Hoechst fluorescence intensity ratio. The mixture contained chlorpyrifos, dimethoate, diazinon, imazalil, iprodione, maneb and mancozeb. HepaRG cells treated with dithiocarbamates were exposed to a mixture containing only maneb and mancozeb. Cells were also treated with a mixture that did not contain maneb and mancozeb (mixture w/o dithiocarbamates). Results are means ± SEM for 5 to 6 independent cultures and are expressed as percentages of the values obtained in untreated -FA-HepaRG cells. *Significantly different from the corresponding untreated or treated -FA-HepaRG cells. #Significantly different from the corresponding untreated -FA or +FA-HepaRG cells (Ctrl).

After 2 weeks, neutral lipids in HepaRG cells cultured with fatty acids (i.e. +FA-HepaRG cells) were moderately but significantly increased as compared to HepaRG cells not exposed to fatty acids (i.e. -FA-HepaRG cells) (Fig. 1b), as previously described (Bucher et al. 2018; Le Guillou et al. 2018). When cells were exposed to the pesticide mixture, neutral lipids were not increased in -FA-HepaRG cells but were enhanced in +FA-HepaRG cells for the 1/5, 1 and 3ADI concentrations (Fig. 1b). The observed effect in +FA-HepaRG cells was also reproduced when these cells were exposed to maneb, mancozeb and maneb + mancozeb (dithiocarbamates) at the 1 and 3ADI concentrations (Fig. 1b). Aggravation of steatosis was also observed when cells were exposed to 2.25 and 6.75 μM MnCl_2_ (Fig. 1b), respectively corresponding to the concentrations of Mn brought with the 1 and 3ADI concentrations of the dithiocarbamates maneb and mancozeb. In contrast, neutral lipids did not significantly increase in +FA-HepaRG cells exposed to a mixture not containing the dithiocarbamates (Fig. 1b), and in +FA-HepaRG cells treated only with diazinon, chlorpyrifos, imazalil, dimethoate, or iprodione (Supplemental Fig. 2). Finally, neutral lipids were not further increased in +FA-HepaRG cells exposed to ethylene thiourea (Supplemental Fig. 2), the main metabolite of the dithiocarbamates maneb and mancozeb (Petitjean et al. 2024).

In MASLD, most neutral lipids accumulating in hepatocytes are made of triglycerides (Ooi et al. 2021; Parlati et al. 2021). Hence, triglycerides were measured in -FA and +FA-HepaRG cells exposed to the pesticide mixture, maneb, mancozeb, or MnCl_2_. In keeping with the Nile red assay, triglycerides were not increased in -FA-HepaRG cells exposed to the pesticide mixture, maneb, mancozeb, or MnCl_2_ (Supplemental Fig. 3). In contrast, exposure to the pesticide mixture, maneb and mancozeb at the 1 and 3ADI concentrations further increased triglyceride accumulation in +FA-HepaRG cells (Supplemental Fig. 3). Exposure of +FA-HepaRG cells to MnCl_2_ significantly augmented triglyceride accumulation only for 6.75 μM, which corresponds to the 3ADI concentration of maneb, or mancozeb (Supplemental Fig. 3).

Altogether, these results showed that chlorpyrifos-ethyl, dimethoate, diazinon, iprodione, imazalil, maneb and mancozeb alone or in a mixture did not induce steatosis in -FA-HepaRG cells after a 2-week exposure. In contrast, maneb, mancozeb or the pesticide mixture worsened the accumulation of total lipids and triglycerides in +FA-HepaRG cells at the 1 and 3 ADI concentrations. Finally, the exacerbation of steatosis was reproduced when +FA-HepaRG cells were exposed to 2.25 and 6.75 μM MnCl_2,_ respectively corresponding to the 1 and 3 ADI concentrations of the dithiocarbamates maneb and mancozeb.

### Effects of maneb, mancozeb and MnCl_2_ in PHH cultured with fatty acids

In a small series of investigations, we wished to determine whether maneb, mancozeb and MnCl_2_ could also aggravate neutral lipid accumulation in PHH cultured in the presence of stearic and oleic acids (+FA-PHH). To this end, +FA-PHH were treated at the ADI concentration for maneb and mancozeb, or with 2.25 μM MnCl_2_, which corresponds to the dithiocarbamates ADI. Of note, these experiments did not include PHH not exposed to stearic and oleic acids because of the low number of PHH that could be obtained. Our results for two different batches of PHH (PHH 1 and 2) confirmed that maneb, mancozeb and MnCl_2_ worsen neutral lipid accumulation in +FA-cells, although the effects of these molecules differed between PHH 1 and 2 (Fig. 2).

**Figure 2.**
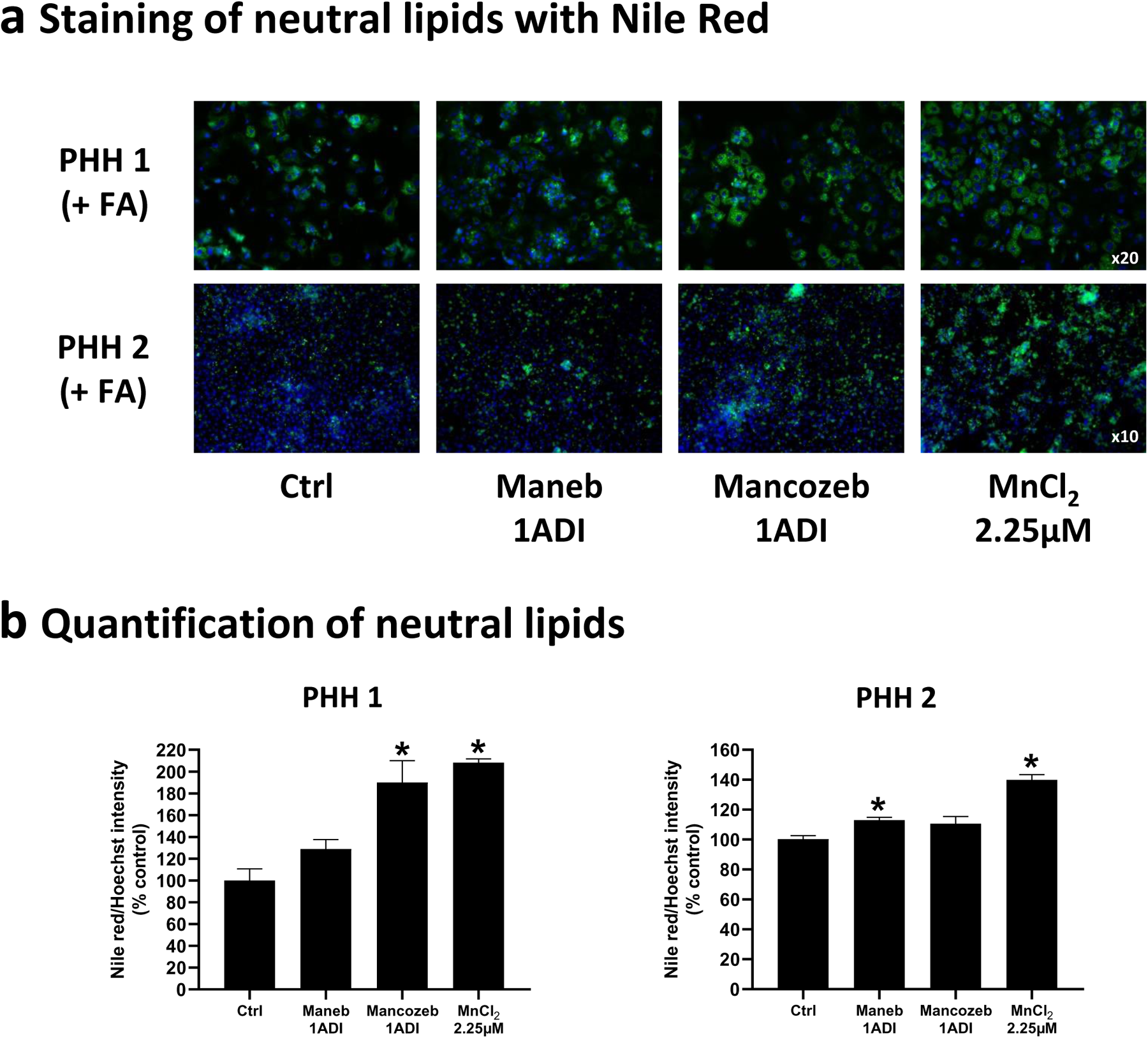
Effects of maneb, mancozeb and manganese chloride on neutral lipid accumulation in two different batches of primary human hepatocytes cultured with stearic and oleic acids. Primary human hepatocytes (PHH) cultured with fatty acids (+FA) were treated or not at the ADI concentration for maneb and mancozeb, or with 2.25 μM manganese chloride (MnCl_2_), which corresponds to the ADI of the dithiocarbamates. Neutral lipids and nuclei were stained with the fluorescent dyes Nile red and Hoechst 33342, respectively. (**a**) Representative pictures of +FA-PHH exposed or not to maneb, mancozeb, or MnCl_2_. Neutral lipids and nuclei are visualized in green and blue, respectively. Pictures were taken at a magnification of ×20 or x10, for PHH 1 and PHH 2, respectively. (**b**) Quantification of neutral lipids in +FA-PHH treated with maneb, mancozeb, or MnCl_2_ using the Nile red/Hoechst fluorescence intensity ratio. Results are means ± SEM for 4 to 6 technical replicates and expressed as percentages of the values obtained in untreated +FA-PHH. *Significantly different from untreated +FA-PHH (Ctrl).

### Mechanisms of dithiocarbamate-induced aggravation of steatosis

To determine the mechanisms whereby dithiocarbamates and MnCl_2_ exacerbated steatosis in +FA-HepaRG cells, we investigated the 4 major metabolic pathways that might be potentially involved, namely fatty acid uptake, mitochondrial FAO, DNL and VLDL secretion, which was assessed via the measurement of apoB in HepaRG cells supernatants.

Fatty acid uptake was not altered by the mixture, maneb and MnCl_2_ after a 24-h treatment (Fig. 3a). However, it was increased by all treatments in both -FA and +FA-HepaRG cells after 48 h although this effect was statistically significant only in +FA-HepaRG cells treated with the mixture or maneb at the ADI concentration (Fig. 3a). We also found that the mixture and maneb profoundly reduced VLDL secretion in both -FA and +FA-HepaRG cells after 24 and 48 h of treatment (Fig. 3b). This effect was reproduced by 2.25 μM MnCl_2_, corresponding to the ADI concentration of maneb.

**Figure 3.**
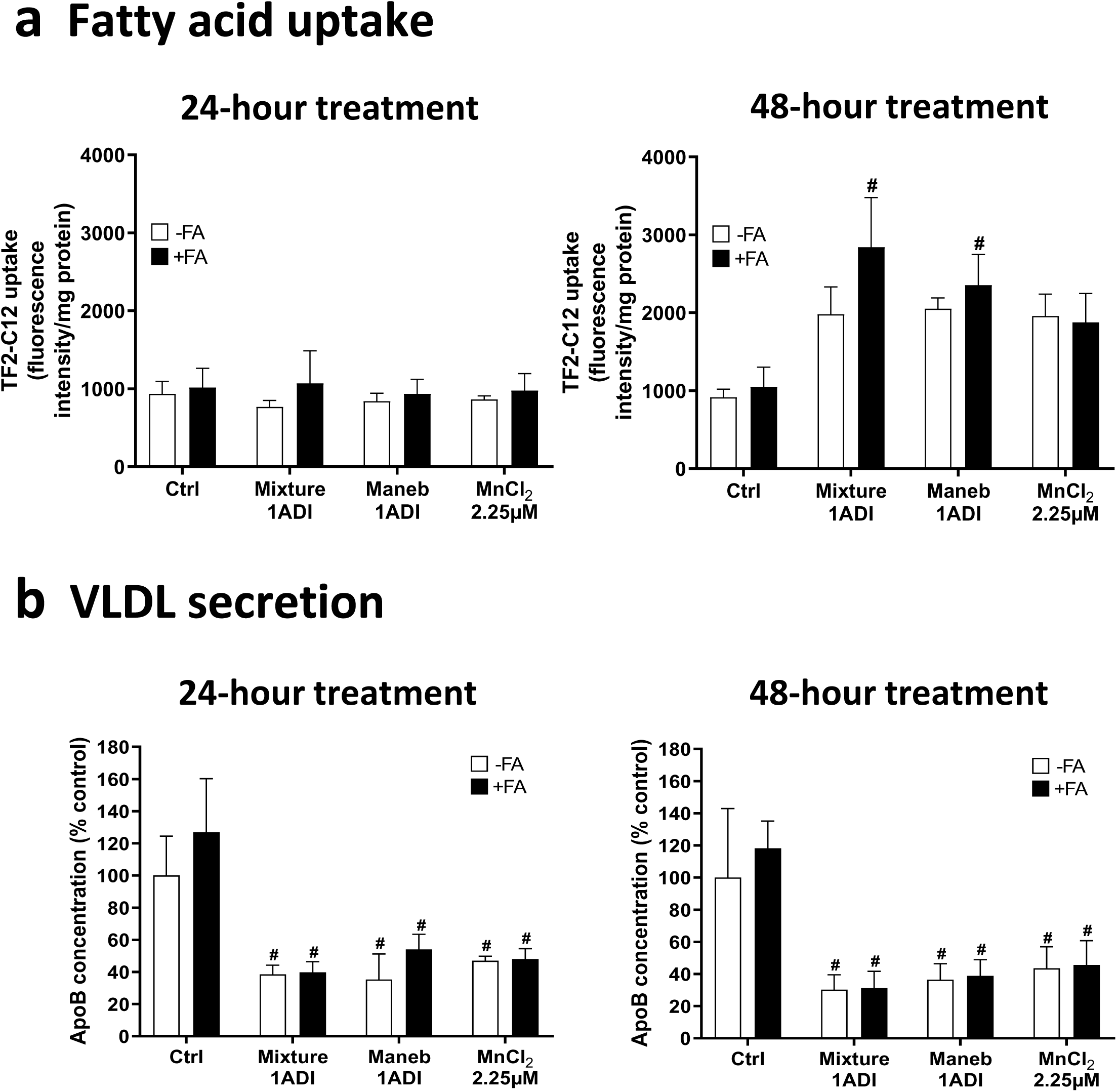
Effects of the pesticide mixture, maneb and manganese chloride on fatty acid uptake and VLDL secretion in differentiated HepaRG cells cultured with or without stearic and oleic acids. HepaRG cells were treated with (+FA) or without (-FA) fatty acids, pesticides, or manganese chloride (MnCl_2_) for 24 or 48 h. Cells were treated at the ADI concentration for the pesticide mixture and maneb, or with 2.25 μM MnCl_2_, which corresponds to the ADI of maneb. (**a**) Fatty acid uptake was measured using the fluorescent probe TF2-C12 after 24 or 48 h of treatment. Results are means ± SEM for 3 independent cultures and are expressed as fluorescence intensity/mg protein. #Significantly different from the corresponding untreated +FA-HepaRG cells (Ctrl). (**b**) VLDL secretion was assessed by apoB measurement in the culture medium after 24 or 48 h of treatment. Results are means ± SEM for 3 to 6 independent cultures and are expressed as percentages of the values obtained in untreated -FA-HepaRG cells. #Significantly different from the corresponding untreated -FA and +FA-HepaRG cells (Ctrl).

Mitochondrial FAO was significantly reduced in +FA-HepaRG cells treated for 2 weeks by the pesticide mixture but only at the 3 ADI concentration (Supplemental Fig. 4). In addition, this metabolic pathway was not altered in -FA and +FA-HepaRG cells treated with different concentrations of maneb, mancozeb, or MnCl_2_ (Supplemental Fig. 4). DNL was significantly reduced in +FA-HepaRG cells compared to -FA-HepaRG cells (Supplemental Fig. 5), confirming that this metabolic pathway is strongly inhibited when fatty acids or lipids are provided in excess (Allard et al. 2021; Ren et al. 2012). DNL was also reduced in -FA-HepaRG cells treated with the mixture, maneb, mancozeb, or MnCl_2_ but these treatments did not alter DNL in +FA-HepaRG cells (Supplemental Fig. 5).

Altogether, these results suggested that the pesticide mixture, maneb and MnCl_2_ could exacerbate steatosis in +FA-HepaRG cells by rapidly impairing VLDL secretion and enhancing fatty acid uptake. Of note, the increase (although not significant) in fatty acid uptake in -FA-HepaRG cells treated with the mixture, maneb, or MnCl_2_ (Fig. 3a) appeared to have no impact on neutral lipids in -FA-HepaRG cells (Fig. 1b). This was most probably because these cells were cultured in the absence of stearic and oleic acids.

### Effect of ZnCl_2_ supplementation on neutral lipids, VLDL secretion and fatty acid uptake

In our recent study, maneb-, mancozeb- and MnCl_2_-induced acute cytotoxicity in differentiated HepaRG cells was significantly prevented by ZnCl_2_ supplementation (Petitjean et al. 2024). Importantly, ZnCl_2_ supplementation reduced (although not significantly) the intracellular levels of MnCl_2_ and significantly restored the depletion of cellular Zn induced by maneb and MnCl_2_ (Petitjean et al. 2024). Hence, in this study, we wished to determine whether Zn supplementation could also prevent the exacerbation of steatosis in +FA-HepaRG cells induced by a 2-week treatment of mancozeb and MnCl_2_. Both 22.5 and 45 μM ZnCl_2_ significantly alleviated the accumulation of neutral lipids in +FA-HepaRG cells treated with mancozeb at the ADI concentration or with 2.25 μM MnCl_2_, corresponding to the ADI of mancozeb (Fig. 4a). Moreover, 45 μM ZnCl_2_ partially alleviated the reduction of VLDL secretion induced by maneb, mancozeb and MnCl_2_ (Fig. 4b), and fully prevented the increase in fatty acid uptake induced by these molecules in +FA-HepaRG cells after 48 h of treatment (Fig. 4c). Altogether, these results suggested that maneb, mancozeb and MnCl_2_ could worsen steatosis and disturb lipid metabolism in +FA-HepaRG cells by disrupting Mn and Zn homeostasis.

**Figure 4.**
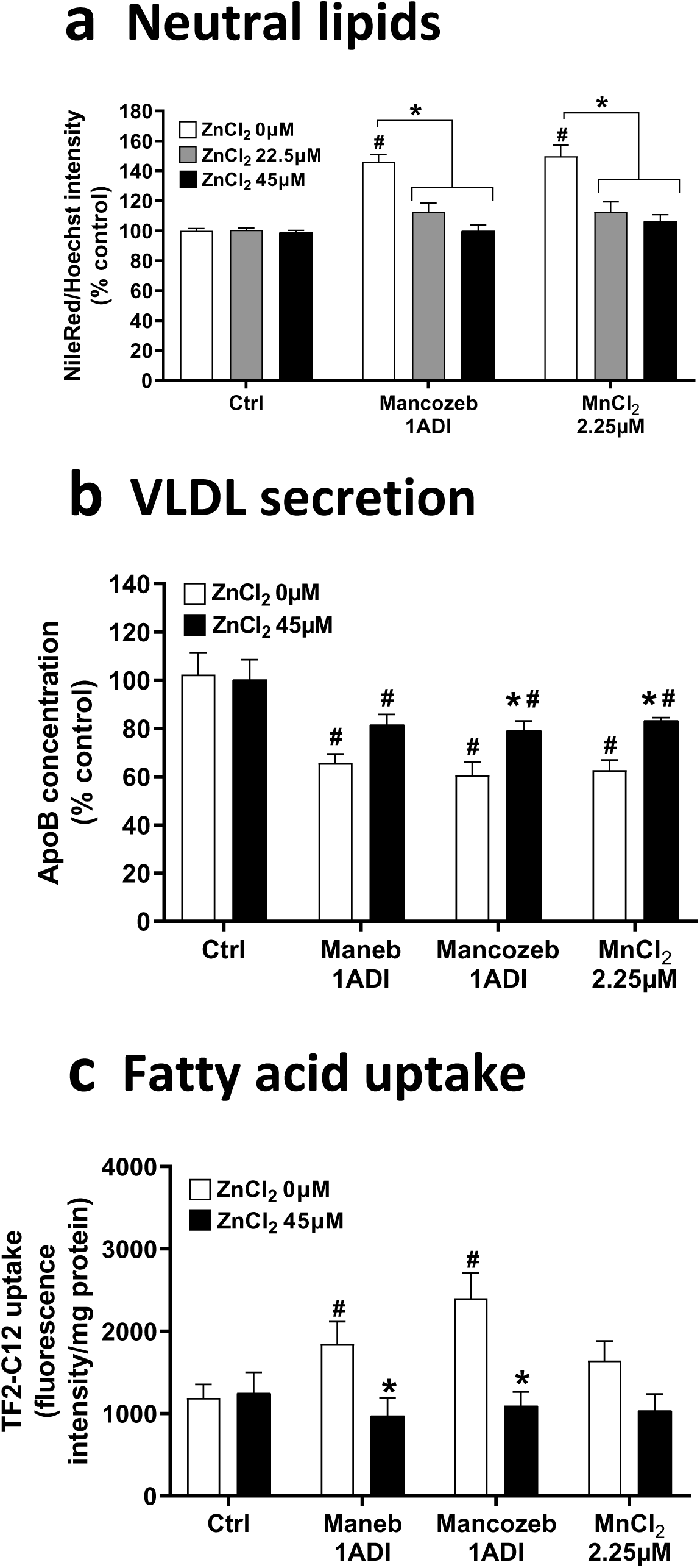
Effects of zinc supplementation on maneb, mancozeb and manganese chloride-induced changes in neutral lipids, VLDL secretion and fatty acid uptake in differentiated HepaRG cells cultured with stearic and oleic acids. HepaRG cells cultured with fatty acids (+FA-HepaRG cells) were treated with or without zinc chloride (ZnCl_2_), mancozeb, maneb, or manganese chloride (MnCl_2_) for 48 h or 2 weeks. Cells were treated at the ADI concentration for mancozeb and maneb, or with 2.25 μM MnCl_2_, which corresponds to the ADI of these dithiocarbamates. Zinc supplementation was achieved with 22.5 or 45 μM ZnCl_2_. (**a**) Neutral lipids. Neutral lipids were assessed after 2 weeks of treatment using the Nile red/Hoechst fluorescence intensity ratio. Results are expressed as percentages of the values obtained in untreated +FA-HepaRG cells. (**b**) VLDL secretion. VLDL secretion was assessed by apoB measurement in the culture medium after 48 h of treatment. Results are expressed as percentages of the values obtained in untreated +FA-HepaRG cells. (**c**) Fatty acid uptake. Fatty acid uptake was measured using the fluorescent probe TF2-C12 after 48 h of treatment. Results are expressed as fluorescence intensity/mg protein. Results for (**a**), (**b**) and (**c**) are means ± SEM for 3 to 4 independent experiments. *Significantly different from the corresponding +FA-HepaRG cells not treated with ZnCl_2_. #Significantly different from the corresponding +FA-HepaRG cells unexposed to dithiocarbamates or MnCl_2_ (Ctrl).

### Expression of genes involved in VLDL secretion and fatty acid uptake

In the next series of investigations, the mRNA expression of different genes involved in VLDL secretion and fatty acid uptake was measured in -FA and +FA-HepaRG cells treated for 24 and 48 h with the pesticide mixture or maneb at the ADI concentration, or with 2.25 μM MnCl_2_, corresponding to the ADI of maneb. Regarding genes involved in VLDL secretion, mRNA levels of apolipoprotein B (*APOB*), apolipoprotein C3 (*APOC3*) and microsomal triglyceride transfer protein (*MTTP*) were reduced by the different treatments after 48 h in -FA and +FA-HepaRG cells, although to different extents (Fig. 5a). However, *APOB* and *APOC3* expression were less or not affected after 24 h (Fig. 5a). Because reduced VLDL secretion can be associated with endoplasmic reticulum (ER) stress (Allard et al. 2021; Ota et al. 2008), mRNA levels of *DDIT3* (*CHOP*), *ERN1* (*IRE1a*), *EIF2AK3* (*PERK*), *HSPA5* (*BIP*) and *ATF6* were measured in -FA and +FA-HepaRG cells treated for 24 and 48 h with the pesticide mixture or maneb at the ADI concentration, or with 2.25 μM MnCl_2_, corresponding to the ADI of maneb. However, the mRNA expression of these different genes was not significantly increased after 24 and 48 h (data not shown), thus suggesting that ER stress was not involved in the impairment of VLDL secretion induced by the pesticide mixture, maneb and MnCl_2_ (Fig. 3a).

**Figure 5.**
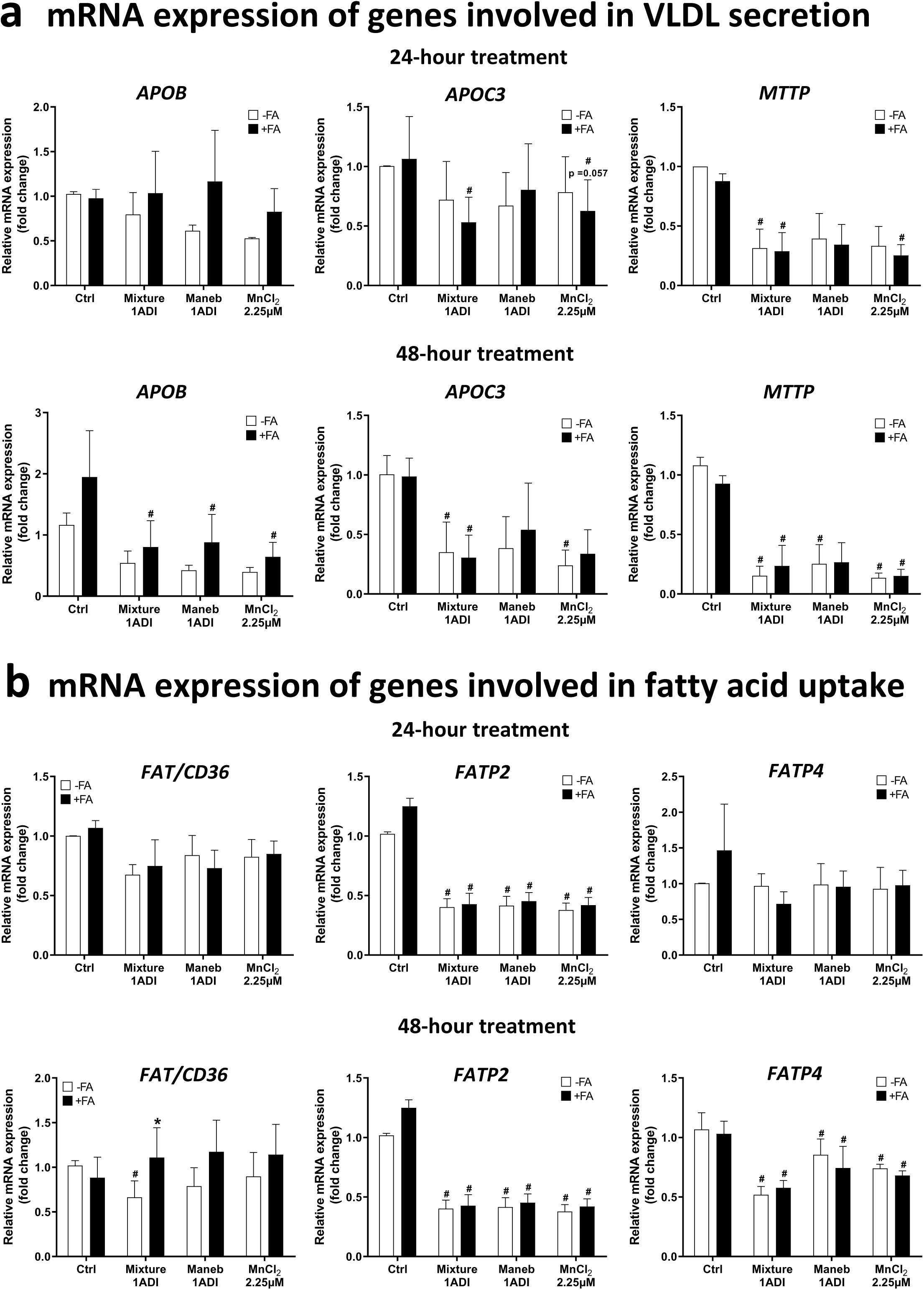
Effects of the pesticide mixture, maneb and manganese chloride on the mRNA expression of genes involved in VLDL secretion and fatty acid uptake in differentiated HepaRG cells cultured with or without stearic and oleic acids. HepaRG cells were treated with (+FA) or without (-FA) fatty acids, pesticides, or manganese chloride (MnCl_2_) for 24 or 48 h. Cells were treated at the ADI concentration for the mixture and maneb, or with 2.25 μM MnCl_2_, which corresponds to the ADI of maneb. (**a**) Expression of *APOB*, *APOC3* and *MTTP* after 24 and 48 h of treatment. (**b**) Expression of *FAT/CD36*, *FATP2* and *FATP4* after 24 and 48 h of treatment. Results for (**a**) and (**b**) are means ± SEM for 3 independent cultures and are expressed as fold change of untreated -FA-HepaRG cells. *Significantly different from the corresponding treated -FA-HepaRG cells. #Significantly different from the corresponding untreated -FA or +FA-HepaRG cells (Ctrl).

Regarding genes involved in fatty acid uptake, mRNA levels of CD36 molecule (fatty acid transporter, *FAT/CD36*), fatty acid transport protein 2 (*FATP2*) and fatty acid transport protein 4 (*FATP4*) were either reduced or unchanged by the different treatments after 24 and 48 h in -FA and +FA-HepaRG cells (Fig. 5b). Hence, these results suggested that higher fatty acid uptake in HepaRG cells treated for 48 h with the pesticide mixture, maneb, or MnCl_2_ (Fig. 3a) could not be explained by the observed changes in the expression of these key fatty acid transporters in hepatocytes.

### Activity and expression of phase I and II XMEs

In the next series of investigations, the activity and expression of several XMEs were measured for two reasons. First, numerous xenobiotics including pesticides have been shown to modulate the expression and/or activity of many XMEs (Abass et al. 2012; Amacher 2010; Hakkola et al. 2020). Second, MASLD has been consistently associated with altered expression and/or activity of different CYPs, UGTs and SULTs (Brill et al. 2012; Cobina and Akhlagi 2017; Smit et al. 2018), as previously mentioned. Notably, preliminary investigations in -FA and +FA-HepaRG cells showed that the pesticide mixture and maneb significantly reduced XME activity at concentrations much lower than 1ADI with reproducible effects for the 1/10ADI concentration. Hence, this low concentration was kept for the remaining investigations reported below.

Regarding the exposure to fatty acids alone, our data showed that CYP2E1 activity and mRNA expression were increased in +FA-HepaRG cells (Fig. 6a), whereas CYP3A4 activity and mRNA expression were reduced in these cells (Fig. 6b), which was fully in line with previous data in experimental and human MASLD (Begriche et al. 2023; Brill et al. 2012; Bucher et al. 2018; Cobina and Akhlagi 2017; Smit et al. 2018). In contrast, the activity of CYP2B6 (Fig. 6c) and that of CYP1A2, CYP2C19, UGT and SULT (Supplemental Fig. 6) were unchanged in +FA-HepaRG cells as compared with -FA-HepaRG cells.

**Figure 6.**
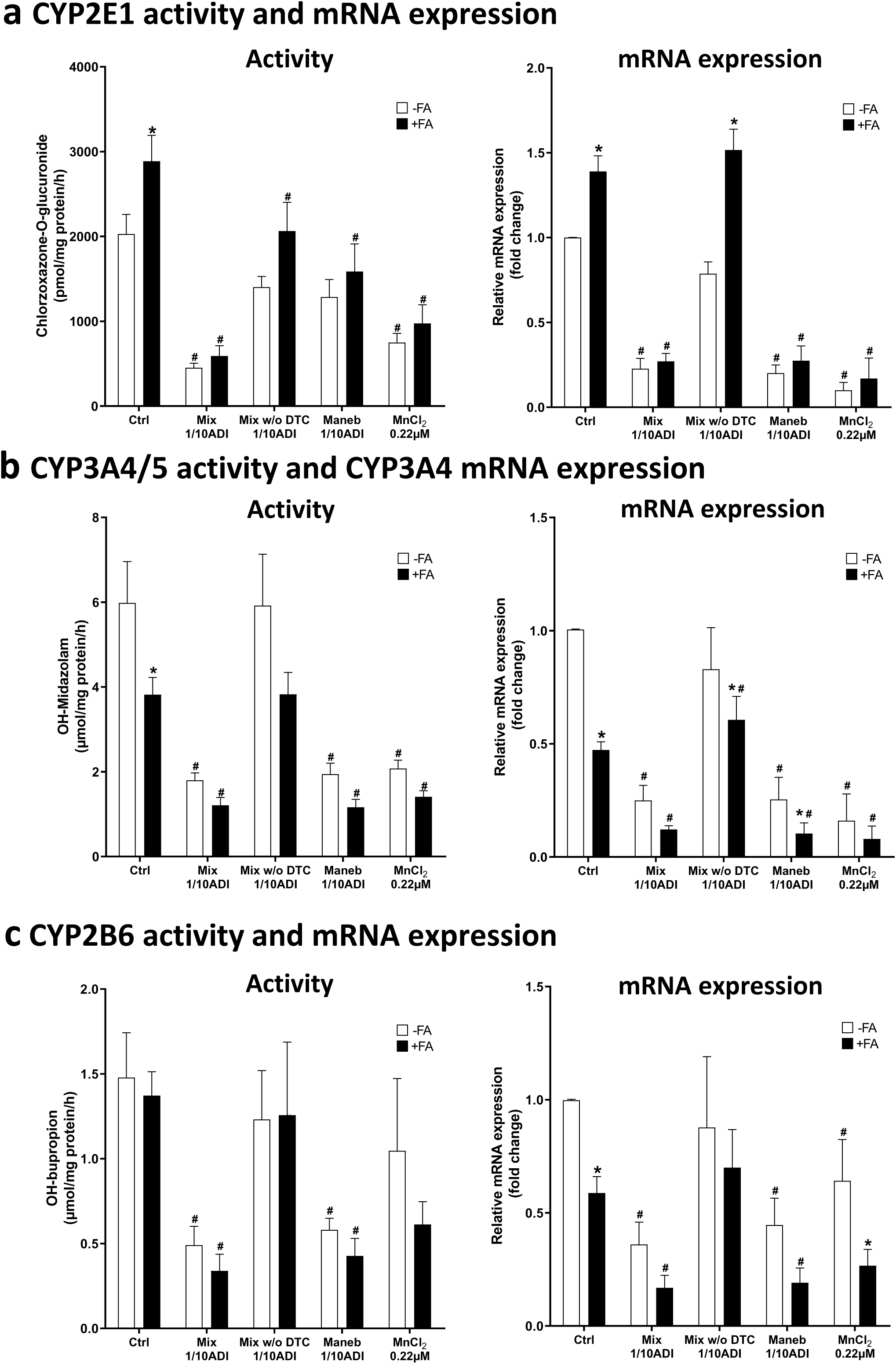
Effects of pesticides and manganese chloride on the mRNA expression and activity of CYP2E1, CYP3A4/5 and CYP2B6 in differentiated HepaRG cells cultured with (+FA) or without (- FA) stearic and oleic acids. HepaRG cells were treated with or without fatty acids, pesticides, or manganese chloride (MnCl_2_) for 2 weeks. Cells were treated at the 1/10 ADI concentration for the pesticide mixture (Mix), the mixture without dithiocarbamates (Mix w/o DTC) and maneb, or with 0.22 μM MnCl_2_, which corresponds to 1/10 ADI of maneb. (**a**) CYP2E1 activity and mRNA expression. CYP2E1 activity, assessed by measuring the formation of chlorzoxazone-O-glucuronide, is expressed in pmol/mg protein/h. *CYP2E1* expression is expressed as fold change of untreated -FA-HepaRG cells. (**b**) CYP3A4 activity and mRNA expression. CYP3A4 activity, assessed by the formation of hydroxymidazolam, is expressed in μmol/mg protein/h. *CYP3A4* expression is expressed as fold change of untreated -FA-HepaRG cells. (**c**) CYP2B6 activity and mRNA expression. CYP2B6 activity, assessed by the formation of hydroxybupropion, is expressed in μmol/mg protein/h. *CYP2B6* expression is expressed as fold change of untreated -FA-HepaRG cells. Results for (**a**), (**b**) and (**c**) are means ± SEM for 3 to 7 independent cultures. *Significantly different from the corresponding untreated or treated -FA-HepaRG cells. #Significantly different from the corresponding untreated -FA or +FA-HepaRG cells (Ctrl).

Concerning pesticide and MnCl_2_ exposure, a reduction of the activity and mRNA expression of CYP2E1, CYP3A4 and CYP2B6 was found in -FA and +FA-HepaRG cells exposed to the pesticide mixture, maneb or MnCl_2_, although to a different extent depending on the experimental conditions (Fig. 6). Notably, the reduction of CYP3A4 and CYP2B6 activity and expression was more pronounced in +FA-HepaRG cells as compared to -FA-HepaRG cells (Figs. 6b and 6c). In contrast, the activity and mRNA expression of CYP2E1, CYP3A4 and CYP2B6 were less or not altered with a mixture not containing the dithiocarbamates maneb and mancozeb (Fig. 6). Finally, the pesticide mixture, maneb and MnCl_2_ did not decrease the activity of CYP1A2, UGT and SULT (Supplemental Fig. 6). However, a significant reduction in CYP2C19 activity was observed only in +FA-HepaRG cells incubated with the pesticide mixture, maneb, or the pesticide mixture not containing the dithiocarbamates (Supplemental Fig. 6). Altogether, these data showed that the pesticide mixture, maneb and MnCl_2_ reduced the expression and activity of CYP2E1, CYP3A4 and CYP2B6, but not that of CYP1A2, UGT and SULT. The decreased activity of CYP2E1, CYP3A4 and CYP2B6 could be due to a transcriptional or post-transcriptional mechanism.

### Expression of genes encoding nuclear receptors

The nuclear receptors AhR, CAR, PXR and farnesoid X-activated receptor (FXR) play a major role in the regulation of the expression of many XMEs including CYPs (Riddick et al. 2023; Toporova and Balaguer 2020). In addition, there is growing evidence that these nuclear receptors also play a role in lipid metabolism (Cai et al. 2021; Toporova and Balaguer 2020; Wang et al. 2022) and MASLD pathophysiology (Cave et al. 2016; Klaunig et al. 2018; Yang et al. 2020). Hence, we assessed the mRNA expression of *AHR*, *CAR*, *FXR* and *PXR* in -FA and +FA-HepaRG cells treated with the pesticide mixture, the mixture without maneb and mancozeb, maneb, or MnCl_2_ for 2 weeks. Overall, these treatments reduced the mRNA levels of *CAR*, *FXR* and *PXR*, although to a different extent depending on the experimental conditions (Fig. 7). In contrast, *AHR* expression was not altered regardless of the experimental conditions (Fig. 7).

**Figure 7.**
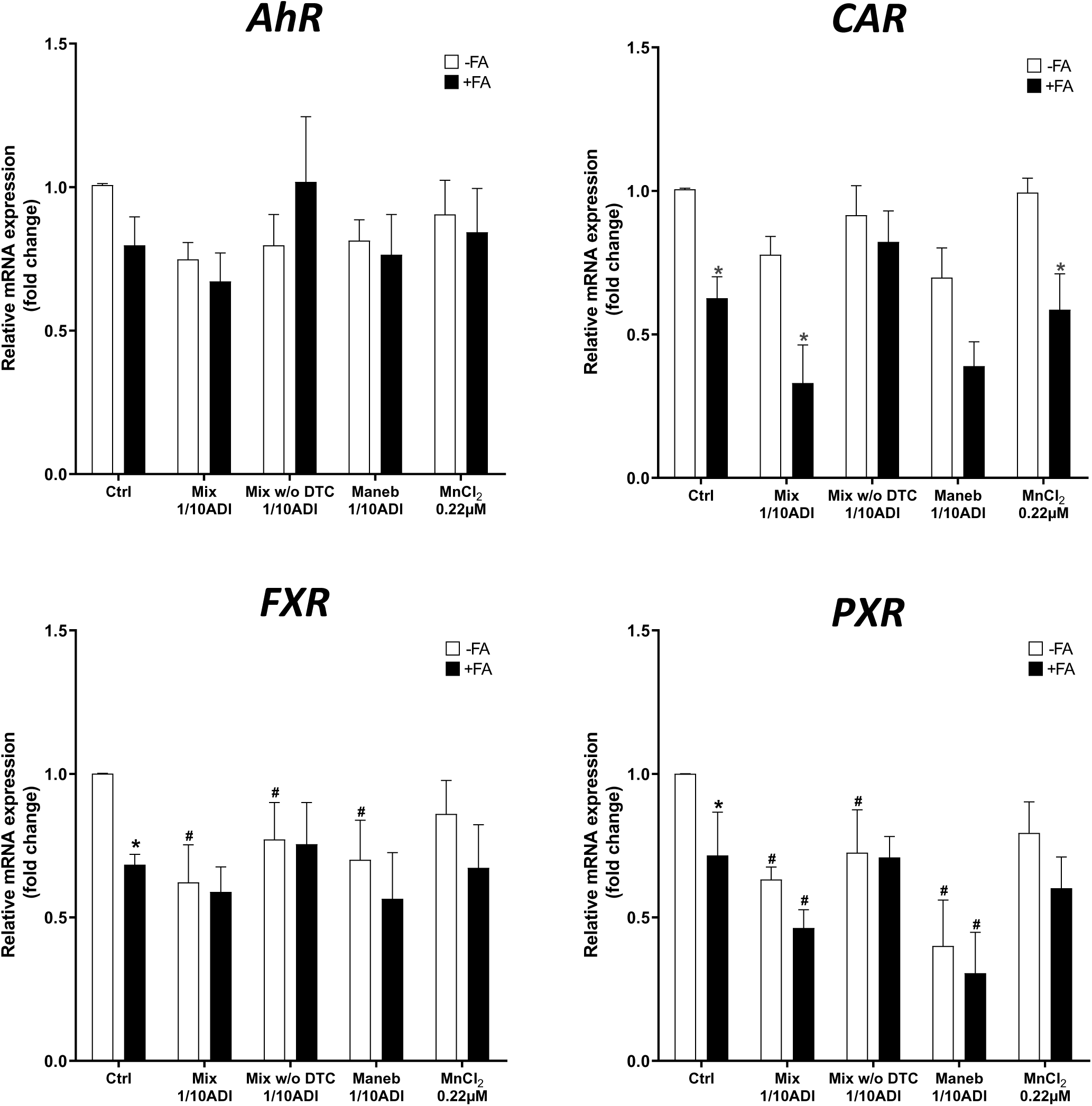
Effects of pesticides and manganese chloride on the mRNA expression of *AHR*, *CAR*, *FXR* and *PXR* in differentiated HepaRG cells cultured with or without stearic and oleic acids. HepaRG cells were treated with (+FA) or without (-FA) fatty acids, pesticides, or manganese chloride (MnCl_2_) for 2 weeks. Cells were treated at the 1/10 ADI concentration for the pesticide mixture (Mix), the mixture without dithiocarbamates (Mix w/o DTC) and maneb, or with 0.22 μM MnCl_2_, which corresponds to 1/10 ADI of maneb. Results are means ± SEM for 3 independent cultures and expressed as fold change of untreated -FA-HepaRG cells. *Significantly different from the corresponding untreated or treated - FA-HepaRG cells. #Significantly different from the corresponding untreated -FA or +FA-HepaRG cells (Ctrl).

### Investigations on diazinon and chlorpyrifos metabolism

In a last series of experiments, we wished to determine whether the pesticide mixture, maneb and MnCl_2_ could alter the metabolism of the organophosphorothioate insecticides diazinon and chlorpyrifos in -FA and +FA-HepaRG cells since reduced activity of CYP3A4 (Fig. 6b), CYP2B6 (Fig. 6c) and CYP2C19 (Supplemental Fig. 6) was found after a 2-week exposure to these treatments. Indeed, diazinon is metabolized to diazoxon and 2-isopropyl-6-methyl-4-pyrimidinol (IMP) by CYP2B6 and CYP3A4 but also by CYP1A2 and CYP2C19 (Kappers et al. 2001; Mutch and Williams 2006; Sams et al. 2004) (Fig. 8a), whereas chlorpyrifos is metabolized to 3,5,6-trichloro-2-pyridinol (TCP) and chlorpyrifos oxon also by these different CYPs (Sams et al. 2004; Tang et al. 2001). Hence, diazinon metabolism was assessed in this study after a 2-week exposure of the HepaRG cells to the pesticide mixture, maneb and MnCl_2_ by measuring the disappearance of diazinon and the production of IMP after an 8- and a 24-h incubation with 50 μM diazinon. Chlorpyrifos metabolism was studied via the production of TCP from 50 μM chlorpyrifos after the same incubation times.

**Figure 8.**
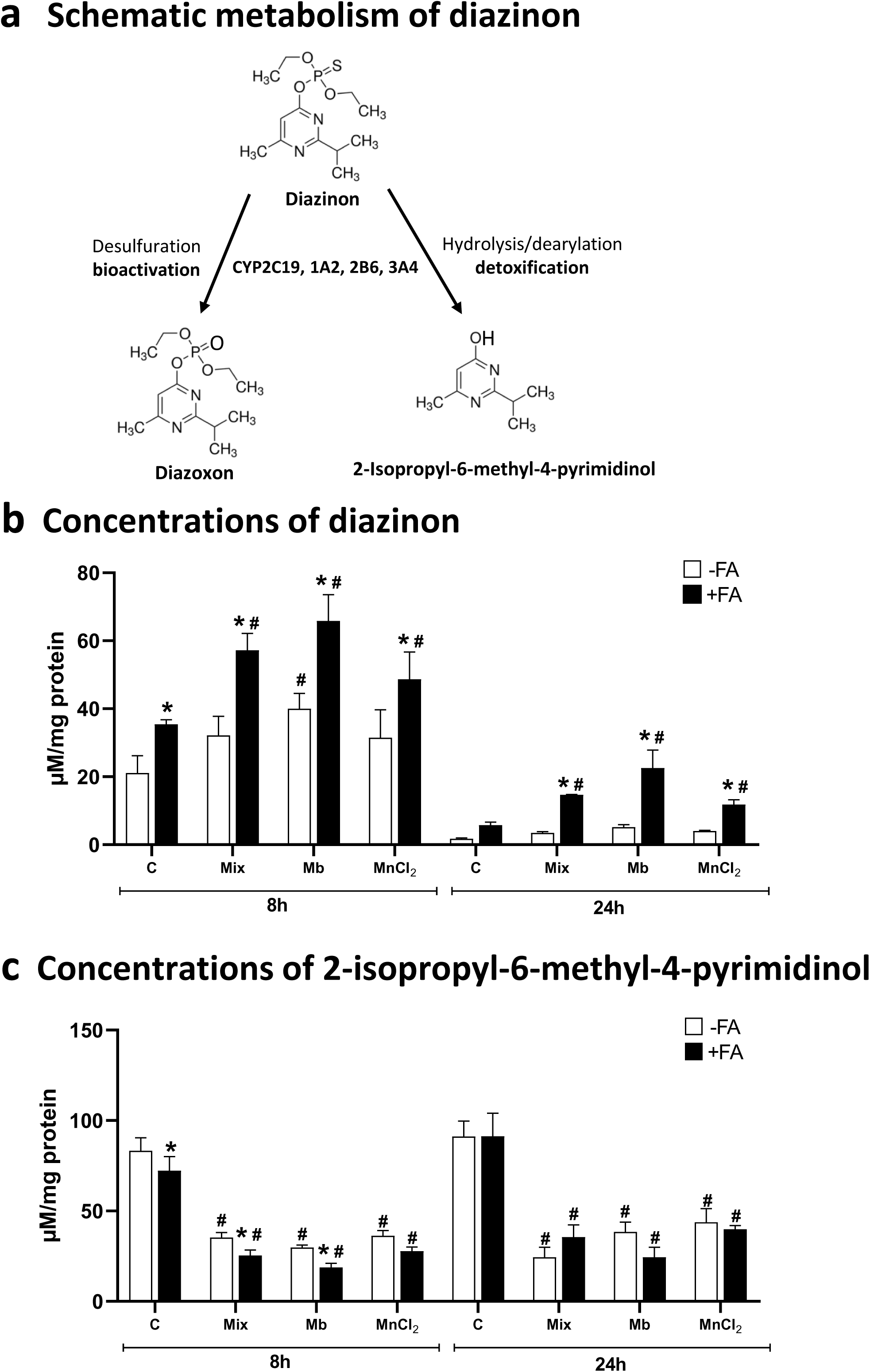
Effects of pesticides and manganese chloride on diazinon metabolism in differentiated HepaRG cells cultured with or without stearic and oleic acids. HepaRG cells were treated with (+FA) or without (-FA) fatty acids, pesticides, or manganese chloride (MnCl_2_) for 2 weeks. Cells were treated at the 1/10 ADI concentration for the pesticide mixture (Mix) and maneb (Mb), or with 0.22 μM MnCl_2_, which corresponds to 1/10 ADI of maneb. (**a**) Schematic metabolism of diazinon. (**b**) and **(c)** Concentrations of diazinon and 2-isopropyl-6-methyl-4-pyrimidinol, respectively. After the 2 weeks, HepaRG cells were treated with 50 µM diazinon and concentrations of remaining diazinon and produced 2-isopropyl-6-methyl-4-pyrimidinol were measured in the culture media after 8 and 24 h. Results for (**b**) and (**c**) are means ± SEM for 4 independent cultures and expressed in μM/mg protein. *Significantly different from the corresponding untreated or treated -FA-HepaRG cells. #Significantly different from the corresponding untreated -FA or +FA-HepaRG cells (C).

Overall, our results showed that the pesticide mixture, maneb and MnCl_2_ lessened both the disappearance of diazinon and the production of IMP and this effect was overall more pronounced in +FA-HepaRG cells as compared to -FA-HepaRG cells (Figs. 8b and 8c). In contrast, these different treatments did not change TCP production from chlorpyrifos regardless of the incubation time (Supplemental Fig. 7). Hence, these results suggested that decreased activity of CYP3A4 (Fig. 6b), CYP2B6 (Fig. 6c) and CYP2C19 (Supplemental Fig. 6) induced by the pesticide mixture, maneb and MnCl_2_ in -FA and +FA-HepaRG cells was sufficient to alter the biotransformation of diazinon but not that of chlorpyrifos.

## Discussion

This study is the first one to show that the dithiocarbamate pesticides maneb and mancozeb can disturb the metabolism of lipids and xenobiotics in differentiated HepaRG cells treated for 2 weeks, possibly via the intracellular release of Mn. Notably, we recently reported that a single treatment of differentiated HepaRG cells with these pesticides induced ROS overproduction and caspase-dependent apoptosis via their biotransformation and the intracellular release of Mn (Petitjean et al. 2024). In the present study, DMSO concentration was lowered to 1% instead of 2% in our previous investigations (Petitjean et al. 2024). This allowed us to significantly curb dithiocarbamate-induced cytotoxicity in the setting of repeated exposures, possibly by lowering the expression and activity of different XMEs including CYPs (Aninat et al. 2006; Dubois-Pot-Schneider et al. 2022; Klein et al. 2014). Indeed, the loss of cellular ATP in treated HepaRG cells was never above 30% of the control cells, even at the highest concentrations of maneb and mancozeb. Altogether, our results indicate that maneb and mancozeb can have broad hepatocellular effects with dire consequences on metabolism, oxidative stress and cell death. Of note, a maximum of 30% decrease in ATP content in HepaRG cells did not impede the study of lipid accumulation and related mechanisms (Allard et al. 2021; Allard et al. 2024). One key finding in this study was the exacerbation of steatosis in +FA-HepaRG cells by maneb, mancozeb and MnCl_2_, whereas these molecules did not induce neutral lipid accumulation in -FA-HepaRG cells. Furthermore, our study disclosed that steatosis aggravation was associated with increased fatty acid uptake and reduced VLDL secretion whereas mitochondrial FAO and DNL were unchanged. Interestingly, impaired VLDL secretion was associated with a reduction in the expression of *APOB* and *APOC3* (two major apolipoproteins in VLDL) and of *MTTP* (also referred to as *MTP*), a key enzyme for the assembly and secretion of this lipoprotein (Allard et al. 2024; Heeren and Scheja 2021). However, the decrease in VLDL secretion and expression of *APOB*, *APOC3* and *MTTP* were not associated with ER stress, contrary to what occurred with different pharmaceuticals inducing steatosis in HepaRG cells such as 5-fluorouracil, indomethacin, rifampicin, troglitazone (Allard et al. 2021) and busulfan (Allard et al. 2024). Thus, further investigations will be needed to determine the exact mechanism(s) whereby maneb, mancozeb and MnCl_2_ reduce *APOB*, *APOC3* and *MTTP* expression and VLDL secretion in hepatocytes.

A previous study reported that short-term treatment with mancozeb worsened neutral lipid accumulation in HepG2 cells cultured with oleic and linoleic acids (Pirozzi et al. 2016). However, aggravation of steatosis was observed with concentrations of mancozeb leading to more than 60% cell mortality. Thus, it is unclear whether the observed effects on cellular lipids were specific, or the mere consequence of severe mitochondrial dysfunction and other major disturbances of cellular homeostasis. In addition, this earlier study did not investigate the effect of mancozeb on fatty acid uptake, mitochondrial FAO, DNL and VLDL secretion.

In our study, mancozeb and MnCl_2_-induced worsening of steatosis in +FA-HepaRG cells was fully prevented by ZnCl_2_ supplementation. In addition, this supplementation restored the impairment of VLDL secretion and alleviated the increase in fatty acid uptake induced by mancozeb, maneb, or MnCl_2_. Notably, Zn supplementation also reduced ROS overproduction, caspase activation and cytotoxicity induced by a single exposure to high concentrations of dithiocarbamates (3ADI) and MnCl_2_ (7 μM) (Petitjean et al. 2024). Hence, these findings strongly suggest that the dithiocarbamates maneb and mancozeb are toxic for the hepatocytes via disruption of Mn and Zn homeostasis, which can be restored by Zn supplementation. Zn supplementation in individuals exposed to high levels of dithiocarbamates might be an efficient means to prevent their potential hepatotoxicity. Although to the best of our knowledge there are no clinical or epidemiological studies reporting dithiocarbamate-induced liver injury, maneb and mancozeb can cause hepatic damage in rodents in particular via oxidative stress and DNA damage (Aprioku et al. 2023; Ben Amara et al. 2015; Nuchniyom et al. 2023; Saber et al. 2019; Sefi et al. 2019; Zhang et al. 2022b). Interestingly, Zn supplementation was reported to prevent maneb-induced teratogenicity in rats (Larsson et al. 1976) and maneb-induced kidney injury in mice (Sefi et al. 2020), thus suggesting that Zn could prevent dithiocarbamate toxicity in different tissues and organs.

Previous epidemiological studies reported a positive association between blood Mn levels and hepatic steatosis (Liu et al. 2023; Spaur et al. 2022), although other investigations found that high blood Mn contents might be a potential protective factor for NAFLD in males (Zhang et al. 2022a). Unfortunately, these studies were not designed to determine whether Mn exposure could aggravate preexisting fatty liver. Interestingly, investigations in the yellow catfish showed that Mn excess can induce hepatic steatosis possibly via increased DNL and reduced mitochondrial FAO (Zhao et al. 2022; Zhao et al. 2023). Moreover, Zn deficiency in rats was reported to induce steatosis depending on the dietary fat composition (Eder and Kirchgessner 1994; Eder and Kirchgessner 1995; tom Dieck et al. 2005). Mechanistic investigations suggested that hepatic lipid accumulation in Zn-deficient rats could be secondary to increased DNL and reduced mitochondrial FAO (Eder and Kirchgessner 1995; tom Dieck et al. 2005). However, these experimental studies did not investigate the effects of Mn overload and Zn deficiency in the liver of animals fed a high-fat diet. Of note, our investigations in differentiated HepaRG cells showed that MnCl_2_ did not induce steatosis in cells that were not cultured with fatty acids (i.e. -FA-HepaRG cells). In addition, MnCl_2_-induced aggravation of steatosis in +FA-HepaRG cells did not involve DNL and mitochondrial FAO. Altogether, these findings suggest that hepatic steatosis induced by Mn excess and Zn deficiency could be specific to some animal species. Furthermore, the mechanisms whereby Mn overload worsens steatosis in +FA-HepaRG cells might differ from those in which Mn excess leads to hepatic lipid accumulation in non-obese animals.

Previous experimental and clinical studies reported that some xenobiotics can aggravate steatosis in the context of preexisting obesity and MASLD. For instance, this has been observed with different pharmaceuticals including the antibiotic tetracycline, the antidiabetic rosiglitazone, the hemorrheologic agent pentoxifylline and some anti-inflammatory corticosteroids (Allard et al. 2019; Massart et al. 2017; Massart et al. 2022). Environmental toxicants reported to worsen fatty liver in obese rodents include bisphenol A, hexabromocyclododecane, nonylphenol, perchloroethylene, 2,3,7,8[tetrachlorodibenzo[p[dioxin and tetrabromodiphenyl ether (Massart et al. 2022; Yanagisawa et al. 2014; Yang et al. 2019). Thus, the present study allows to add maneb, mancozeb and Mn to the growing number of molecules that could worsen obesity-associated steatosis.

Obesity and MASLD are associated with altered activity of different phase I XMEs including CYPs (Brill et al. 2012; Cobina and Akhlagi 2017; Smit et al. 2018). Accordingly, CYP2E1 activity was increased while CYP3A4/5 activity was decreased in +FA-HepaRG cells in comparison with -FA-HepaRG cells. These results are fully in line with several clinical investigations carried out in obese patients with MASLD (Begriche et al. 2023; Brill et al. 2012; Chalasani et al. 2003; Cobina and Akhlagi 2017; Kolwankar et al. 2007; Smit et al. 2018). In addition to obesity and related metabolic diseases, numerous xenobiotics including pesticides can modulate the expression and activity of many XMEs (Abass et al. 2012; Amacher 2010; Hakkola et al. 2020). In this study, low concentration (1/10 ADI) of maneb significantly reduced the activity of CYP2E1, CYP3A4/5, CYP2B6 and CYP2C19 in - FA and +FA-HepaRG cells. Of note, pesticides can be metabolized by a large repertoire of XMEs. For instance, diazinon and chlorpyrifos are metabolized by different CYPs including CYP1A2, CYP2B6, CYP2C19 and CYP3A4 (Kappers et al. 2001; Mutch and Williams 2006; Sams et al. 2004; Tang et al. 2001).

Hence, there can be numerous interactions between MASLD and pesticide exposure as regards xenobiotic biotransformation. In keeping with this hypothesis, alteration of diazinon biotransformation induced by low concentration of maneb was exacerbated in +FA-HepaRG cells as compared to -FA-HepaRG cells. In contrast, chlorpyrifos biotransformation to 3,5,6-trichloro-2-pyridinol was not impaired by maneb. Although we did not have a definite explanation for this difference, chlorpyrifos metabolism might be less dependent on CYP3A4/5, CYP2B6 and CYP2C19 than is diazinon. Noteworthy, various pharmaceuticals are metabolized by these CYPs (Hedrich et al. 2016; Hirota et al. 2013; Klyushova et al. 2022). Hence, individuals exposed to dithiocarbamates might be at risk for drug-drug interactions and related adverse effects.

CAR and PXR are key nuclear receptors regulating CYP3A4/5 and CYP2B6 expression (Cai et al. 2021; Hedrich et al. 2016; Klyushova et al. 2022). Interestingly, the pesticide mixture, maneb and MnCl_2_ decreased *CAR* and *PXR* mRNA levels in -FA- and +FA-HepaRG cells although to a different extent depending on the experimental conditions. Hence, the decrease in *CAR* and *PXR* mRNA levels in some experimental settings might explain, at least in part, the reduction of *CYP3A4* and *CYP2D6* expression. Of note, there is growing evidence that CAR and PXR also regulate the metabolism of carbohydrates and lipids (Cai et al. 2021; Cave et al. 2016). Thus, further investigations would be required to determine whether dithiocarbamate-induced reduction in *CAR* and *PXR* expression participates in the aggravation of steatosis in +FA-HepaRG cells. The pesticide mixture and maneb also decreased the expression of *FXR*, which plays a major role in bile acid metabolism (Sun et al. 2021). Of note, drug-induced downregulation of FXR signaling favors intrahepatic cholestasis (Ding et al. 2018; Petrov et al. 2021). Hence, individuals exposed to dithiocarbamates might also be at risk for cholestatic liver disease.

In conclusion, this study showed that maneb, mancozeb and MnCl_2_ aggravated lipid accumulation in HepaRG cells cultured with stearic and oleic acids. In contrast, chlorpyrifos, dimethoate, diazinon, iprodione and imazalil did not have such an effect. Moreover, low concentrations of maneb and MnCl_2_ altered the expression and activity of different CYPs in +FA- and -FA-HepaRG cells, which was associated with an alteration of diazinon biotransformation. These findings could have major pathophysiological consequences in obese individuals with MASLD and exposed to the dithiocarbamates maneb and mancozeb. In addition, our investigations suggested that Zn supplementation might be an efficient strategy to prevent dithiocarbamate-induced disturbances of liver metabolism in the context of MASLD.

## Acknowledgments

We are grateful to INSERM (Institut National de la Santé et de la Recherche Médicale), CNRS (Centre National de la Recherche Scientifique) and the University of Rennes for their constant financial support. We would like to thank Dr Kyle Hoehn (University of New South Wales, Sydney, Australia) for having given us the detailed protocol to investigate *de novo* lipogenesis in cultured cells. We also thank Korydwen Fruit for her assistance in some investigations.

## Authors’ contributions

Conception and design: Camille C. Savary, Anne Corlu, Pascal Loyer, and Bernard Fromenty. Methodology: Kilian Petitjean, Laurence Amalric, Nicole Baran, Camille Savary, Pascal Loyer, and Bernard Fromenty. Material preparation, data collection, and analysis: Kilian Petitjean, Giovanna Dicara, Sébastien Bristeau, and Hugo Coppens-Exandier. Statistical analysis: Kilian Petitjean. Supervision: Anne Corlu, Pascal Loyer, and Bernard Fromenty. The first draft of the manuscript was written by Kilian Petitjean and Bernard Fromenty and all authors commented on previous versions of the manuscript. All authors read and approved the final manuscript.

## Data availability

The authors declare that the data supporting the findings of this study are available within the paper and its Supplemental Figures. Should any raw data files be needed in another format they are available from the corresponding author upon reasonable request.

## Declarations

### Ethics approval

Not applicable.

## References

Abass K, Lämsä V, Reponen P, Küblbeck J, Honkakoski P, Mattila S, et al. Characterization of human cytochrome P450 induction by pesticides. Toxicology. 2012;294(1):17–26.

Allard J, Le Guillou D, Begriche K, Fromenty B. Drug-induced liver injury in obesity and nonalcoholic fatty liver disease. Adv Pharmacol. 2019;85:75–107.

Allard J, Bucher S, Massart J, Ferron PJ, Le Guillou D, Loyant R, et al. Drug-induced hepatic steatosis in absence of severe mitochondrial dysfunction in HepaRG cells: proof of multiple mechanism-based toxicity. Cell Biol Toxicol. 2021;37(2):151–75.

Allard J, Bucher S, Ferron PJ, Launay Y, Fromenty B. Busulfan induces steatosis in HepaRG cells but not in primary human hepatocytes: Possible explanations and implication for the prediction of drug-induced liver injury. Fundam Clin Pharmacol. 2024;38(1):152–167.

Amacher DE. The effects of cytochrome P450 induction by xenobiotics on endobiotic metabolism in pre-clinical safety studies. Toxicol Mech Methods. 2010;20(4):159–66.

Aninat C, Piton A, Glaise D, Le Charpentier T, Langouët S, Morel F, et al. Expression of cytochromes P450, conjugating enzymes and nuclear receptors in human hepatoma HepaRG cells. Drug Metab Dispos. 2006;34(1):75–83.

Anthérieu S, Chesné C, Li R, Camus S, Lahoz A, Picazo L, et al. Stable expression, activity, and inducibility of cytochromes P450 in differentiated HepaRG cells. Drug Metab Dispos. 2010;38(3):516–25.

Aprioku JS, Amamina AM, Nnabuenyi PA. Mancozeb-induced hepatotoxicity: protective role of curcumin in rat animal model. Toxicol Res (Camb). 2023;12(1):107–116.

Arab JP, Addolorato G, Mathurin P, Thursz MR. Alcohol-Associated Liver Disease: Integrated Management With Alcohol Use Disorder. Clin Gastroenterol Hepatol. 2023;21(8):2124–34.

Araújo RAL, Cremonese C, Santos R, Piccoli C, Carvalho G, Freire C, et al. Association of occupational exposure to pesticides with overweight and abdominal obesity in family farmers in southern Brazil. Int J Environ Health Res. 2022;32(12):2798–809.

Aubert J, Begriche K, Knockaert L, Robin MA, Fromenty B. Increased expression of cytochrome P450 2E1 in nonalcoholic fatty liver disease: mechanisms and pathophysiological role. Clin Res Hepatol Gastroenterol. 2011;35(10):630–7.

Begriche K, Penhoat C, Bernabeu-Gentey P, Massart J, Fromenty B. Acetaminophen-induced hepatotoxicity in obesity and nonalcoholic fatty liver disease: a critical review. Livers. 2023;3(1):33–53.

Ben Amara I, Ben Saad H, Hamdaoui L, Karray A, Boudawara T, Ben Ali Y, et al. Maneb disturbs expression of superoxide dismutase and glutathione peroxidase, increases reactive oxygen species production, and induces genotoxicity in liver of adult mice. Environ Sci Pollut Res Int. 2015;22(16):12309–22.

Brill MJ, Diepstraten J, van Rongen A, van Kralingen S, van den Anker JN, Knibbe CA. Impact of obesity on drug metabolism and elimination in adults and children. Clin Pharmacokinet. 2012;51(5):277–304.

Brunt EM, Kleiner DE, Carpenter DH, Rinella M, Harrison SA, Loomba R, et al. NAFLD: Reporting Histologic Findings in Clinical Practice. Hepatology. 2021;73(5):2028–38.

Bucher S, Le Guillou D, Allard J, Pinon G, Begriche K, Tête A, et al. Possible Involvement of Mitochondrial Dysfunction and Oxidative Stress in a Cellular Model of NAFLD Progression Induced by Benzo[a]pyrene/Ethanol CoExposure. Oxid Med Cell Longev. 2018;2018:4396403.

Cai X, Young GM, Xie W. The xenobiotic receptors PXR and CAR in liver physiology, an update. Biochim Biophys Acta Mol Basis Dis. 2021;1867(6):166101.

Caldas ED, Miranda MC, Conceição MH, de Souza LC. Dithiocarbamates residues in Brazilian food and the potential risk for consumers. Food Chem Toxicol. 2004;42(11):1877–83.

Cano L, Desquilles L, Ghukasyan G, Angenard G, Landreau C, Corlu A, et al. SARS-CoV-2 receptor ACE2 is upregulated by fatty acids in human MASH. JHEP Rep. 2023;6(1):100936.

Cave MC, Clair HB, Hardesty JE, Falkner KC, Feng W, Clark BJ, et al. Nuclear receptors and nonalcoholic fatty liver disease. Biochim Biophys Acta. 2016;1859(9):1083–99.

Cerec V, Glaise D, Garnier D, Morosan S, Turlin B, Drenou B, et al. Transdifferentiation of hepatocyte-like cells from the human hepatoma HepaRG cell line through bipotent progenitor. Hepatology. 2007, 45(4):957–67.

Chalasani N, Gorski JC, Asghar MS, Asghar A, Foresman B, Hall SD, et al. Hepatic cytochrome P450 2E1 activity in nondiabetic patients with nonalcoholic steatohepatitis. Hepatology. 2003;37(3):544–50.

Cobbina E, Akhlaghi F. Non-alcoholic fatty liver disease (NAFLD) - pathogenesis, classification, and effect on drug metabolizing enzymes and transporters. Drug Metab Rev. 2017;49(2):197–211.

Ding L, Zhang B, Li J, Yang L, Wang Z. Beneficial effect of resveratrol on α[naphthyl isothiocyanate[induced cholestasis via regulation of the FXR pathway. Mol Med Rep. 2018;17(1):1863–72.

Dubois-Pot-Schneider H, Aninat C, Kattler K, Fekir K, Jarnouen K, Cerec V, et al. Transcriptional and Epigenetic Consequences of DMSO Treatment on HepaRG Cells. Cells. 2022;11(15):2298.

EASL, European Association for the Study of the Liver. EASL Clinical Practice Guideline: Occupational liver diseases. J Hepatol. 2019;71(5):1022–1037.

Eder K, Kirchgessner M. Dietary fat influences the effect of zinc deficiency on liver lipids and fatty acids in rats force-fed equal quantities of diet. J Nutr. 1994;124(10):1917–26.

Eder K, Kirchgessner M. Zinc deficiency and activities of lipogenic and glycolytic enzymes in liver of rats fed coconut oil or linseed oil. Lipids. 1995;30(1):63–9.

EFSA, European Food Safety Authority. Annual report on pesticide residues according to article 32 of regulation (EC) No 396/2005. EFSA J 2010;8(7),1646.

EFSA, European Food Safety Authority. The 2015 European Union report on pesticide residues in food. EFSA J 2017;15(4):e04791.

Evangelou E, Ntritsos G, Chondrogiorgi M, Kavvoura FK, Hernández AF, Ntzani EE, et al. Exposure to pesticides and diabetes: A systematic review and meta-analysis. Environ Int. 2016;91:60–8.

Ferron PJ, Le Daré B, Bronsard J, Steichen C, Babina E, Pelletier R, et al. Molecular Networking for Drug Toxicities Studies: The Case of Hydroxychloroquine in COVID-19 Patients. Int J Mol Sci. 2021;23(1):82.

Fromenty B, Lettéron P, Fisch C, Berson A, Deschamps D, Pessayre D. Evaluation of human blood lymphocytes as a model to study the effects of drugs on human mitochondria. Effects of low concentrations of amiodarone on fatty acid oxidation, ATP levels and cell survival. Biochem Pharmacol. 1993;46(3):421–32.

Fromenty B. Inhibition of mitochondrial fatty acid oxidation in drug-induced hepatic steatosis. Liver Res. 2019;3(3-4),157–69.

Fromenty B, Roden M. Mitochondrial alterations in fatty liver diseases. J Hepatol. 2023;78(2):415–29.

Greenspan P, Mayer EP, Fowler SD. Nile red: a selective fluorescent stain for intracellular lipid droplets. J Cell Biol. 1985;100(3):965–73.

Grünig D, Duthaler U, Krähenbühl S. Effect of Toxicants on Fatty Acid Metabolism in HepG2 Cells. Front Pharmacol. 2018;9:257.

Hakkola J, Hukkanen J, Turpeinen M, Pelkonen O. Inhibition and induction of CYP enzymes in humans: an update. Arch Toxicol. 2020;94(11):3671–722.

Hedrich WD, Hassan HE, Wang H. Insights into CYP2B6-mediated drug-drug interactions. Acta Pharm Sin B. 2016;6(5):413–25.

Heeren J, Scheja L. Metabolic-associated fatty liver disease and lipoprotein metabolism. Mol Metab. 2021;50:101238.

Hirota T, Eguchi S, Ieiri I. Impact of genetic polymorphisms in CYP2C9 and CYP2C19 on the pharmacokinetics of clinically used drugs. Drug Metab Pharmacokinet. 2013;28(1):28–37.

Hochane M, Trichet V, Pecqueur C, Avril P, Oliver L, Denis J, et al. Low-Dose Pesticide Mixture Induces Senescence in Normal Mesenchymal Stem Cells (MSC) and Promotes Tumorigenic Phenotype in Premalignant MSC. Stem Cells. 2017;35(3):800–11.

Kappers WA, Edwards RJ, Murray S, Boobis AR. Diazinon is activated by CYP2C19 in human liver. Toxicol Appl Pharmacol. 2001;177(1):68–76.

Klaunig JE, Li X, Wang Z. Role of xenobiotics in the induction and progression of fatty liver disease. Toxicol Res (Camb). 2018;7(4):664–80.

Klein S, Mueller D, Schevchenko V, Noor F. Long-term maintenance of HepaRG cells in serum-free conditions and application in a repeated dose study. J Appl Toxicol. 2014;34(10):1078–86.

Klyushova LS, Perepechaeva ML, Grishanova AY. The Role of CYP3A in Health and Disease. Biomedicines. 2022;10(11):2686.

Kolwankar D, Vuppalanchi R, Ethell B, Jones DR, Wrighton SA, Hall SD, et al. Association between nonalcoholic hepatic steatosis and hepatic cytochrome P-450 3A activity. Clin Gastroenterol Hepatol. 2007;5(3):388–93.

Lamat H, Sauvant-Rochat MP, Tauveron I, Bagheri R, Ugbolue UC, Maqdasi S, et al. Metabolic syndrome and pesticides: A systematic review and meta-analysis. Environ Pollut. 2022;305:119288.

Larrain S, Rinella ME. A myriad of pathways to NASH. Clin Liver Dis. 2012;16(3):525–48.

Larsson KS, Arnander C, Cekanova E, Kjellberg M. Studies of teratogenic effects of the dithiocarbamates maneb, mancozeb, and propineb. Teratology. 1976;14(2):171–83.

Le Guillou D, Bucher S, Begriche K, Hoët D, Lombès A, Labbe G, et al. Drug-Induced Alterations of Mitochondrial DNA Homeostasis in Steatotic and Nonsteatotic HepaRG Cells. J Pharmacol Exp Ther. 2018;365(3):711–26.

Liu J, Tan L, Liu Z, Shi R. Blood and urine manganese exposure in non-alcoholic fatty liver disease and advanced liver fibrosis: an observational study. Environ Sci Pollut Res Int. 2023;30(9):22222–31.

Massart J, Begriche K, Moreau C, Fromenty B. Role of nonalcoholic fatty liver disease as risk factor for drug-induced hepatotoxicity. J Clin Transl Res. 2017;3(Suppl 1):212–32.

Massart J, Begriche K, Fromenty B. Cytochrome P450 2E1 should not be neglected for acetaminophen-induced liver injury in metabolic diseases with altered insulin levels or glucose homeostasis. Clin Res Hepatol Gastroenterol. 2021;45(1):101470.

Massart J, Begriche K, Corlu A, Fromenty B. Xenobiotic-Induced Aggravation of Metabolic-Associated Fatty Liver Disease. Int J Mol Sci. 2022;23(3):1062.

Mutch E, Williams FM. Diazinon, chlorpyrifos and parathion are metabolised by multiple cytochromes P450 in human liver. Toxicology. 2006;224(1-2):22–32.

Nuchniyom P, Intui K, Laoung-On J, Jaikang C, Quiggins R, Photichai K, et al. Effects of Nelumbonucifera Gaertn. Petal Tea Extract on Hepatotoxicity and Oxidative Stress Induced by Mancozeb in Rat Model. Toxics. 2023;11(6):480.

Ooi GJ, Meikle PJ, Huynh K, Earnest A, Roberts SK, Kemp W, et al. Hepatic lipidomic remodeling in severe obesity manifests with steatosis and does not evolve with non-alcoholic steatohepatitis. J Hepatol. 2021;75(3):524–35.

Ota T, Gayet C, Ginsberg HN. Inhibition of apolipoprotein B100 secretion by lipid-induced hepatic endoplasmic reticulum stress in rodents. J Clin Invest. 2008;118(1):316–32.

Parlati L, Régnier M, Guillou H, Postic C. New targets for NAFLD. JHEP Rep. 2021;3(6):100346.

Petitjean K, Verres Y, Bristeau S, Ribault C, Aninat C, Olivier C, et al. Low concentrations of ethylene bisdithiocarbamate pesticides maneb and mancozeb impair manganese and zinc homeostasis to induce oxidative stress and caspase-dependent apoptosis in human hepatocytes. Chemosphere. 2024;346:140535.

Petrov PD, Soluyanova P, Sánchez-Campos S, Castell JV, Jover R. Molecular mechanisms of hepatotoxic cholestasis by clavulanic acid: Role of NRF2 and FXR pathways. Food Chem Toxicol. 2021;158:112664.

Pirozzi AV, Stellavato A, La Gatta A, Lamberti M, Schiraldi C. Mancozeb, a fungicide routinely used in agriculture, worsens nonalcoholic fatty liver disease in the human HepG2 cell model. Toxicol Lett. 2016;249:1–4.

Quesnot N, Bucher S, Gade C, Vlach M, Vene E, Valença S, et al. Production of chlorzoxazone glucuronides via cytochrome P4502E1 dependent and independent pathways in human hepatocytes. Arch Toxicol. 2018;92(10):3077–91.

Rebouillat P, Vidal R, Cravedi JP, Taupier-Letage B, Debrauwer L, Gamet-Payrastre L, et al. Prospective association between dietary pesticide exposure profiles and type 2 diabetes risk in the NutriNet-Santé cohort. Environ Health. 2022;21(1):57.

Ren LP, Chan SM, Zeng XY, Laybutt DR, Iseli TJ, Sun RQ, et al. Differing endoplasmic reticulum stress response to excess lipogenesis versus lipid oversupply in relation to hepatic steatosis and insulin resistance. PLoS One. 2012;7(2):e30816.

Riddick DS. Fifty Years of Aryl Hydrocarbon Receptor Research as Reflected in the Pages of Drug Metabolism and Disposition. Drug Metab Dispos. 2023;51(6):657–71.

Rinella ME, Lazarus JV, Ratziu V, Francque SM, Sanyal AJ, Kanwal F, et al. A multisociety Delphi consensus statement on new fatty liver disease nomenclature. J Hepatol. 2023;79(6):1542–56.

Saber TM, Abo-Elmaaty AMA, Abdel-Ghany HM. Curcumin mitigates mancozeb-induced hepatotoxicity and genotoxicity in rats. Ecotoxicol Environ Saf. 2019;183:109467.

Sams C, Cocker J, Lennard MS. Biotransformation of chlorpyrifos and diazinon by human liver microsomes and recombinant human cytochrome P450s (CYP). Xenobiotica. 2004;34(10):861–73.

Sang H, Lee KN, Jung CH, Han K, Koh EH. Association between organochlorine pesticides and nonalcoholic fatty liver disease in the National Health and Nutrition Examination Survey 2003-2004. Sci Rep. 2022;12(1):11590.

Sefi M, Elwej A, Chaâbane M, Bejaoui S, Marrekchi R, Jamoussi K, et al. Beneficial role of vanillin, a polyphenolic flavoring agent, on maneb-induced oxidative stress, DNA damage, and liver histological changes in Swiss albino mice. Hum Exp Toxicol. 2019;38(6):619–31.

Sefi M, Chaâbane M, Elwej A, Bejaoui S, Marrekchi R, Jamoussi K, et al. Zinc alleviates maneb-induced kidney injury in adult mice through modulation of oxidative stress, genotoxicity, and histopathological changes. Environ Sci Pollut Res Int. 2020;27(8):8091–102.

Smit C, De Hoogd S, Brüggemann RJM, Knibbe CAJ. Obesity and drug pharmacology: a review of the influence of obesity on pharmacokinetic and pharmacodynamic parameters. Expert Opin Drug Metab Toxicol. 2018;14(3):275–85.

Spaur M, Nigra AE, Sanchez TR, Navas-Acien A, Lazo M, Wu HC. Association of blood manganese, selenium with steatosis, fibrosis in the National Health and Nutrition Examination Survey, 2017-18. Environ Res. 2022;213:113647.

Ssemugabo C, Bradman A, Ssempebwa JC, Sillé F, Guwatudde D. An assessment of health risks posed by consumption of pesticide residues in fruits and vegetables among residents in the Kampala Metropolitan Area in Uganda. Int J Food Contam. 2022;9(1):4.

Sun L, Cai J, Gonzalez FJ. The role of farnesoid X receptor in metabolic diseases, and gastrointestinal and liver cancer. Nat Rev Gastroenterol Hepatol. 2021;18(5):335–47.

Tang J, Cao Y, Rose RL, Brimfield AA, Dai D, Goldstein JA, et al. Metabolism of chlorpyrifos by human cytochrome P450 isoforms and human, mouse, and rat liver microsomes. Drug Metab Dispos. 2001;29(9):1201–4.

tom Dieck H, Döring F, Fuchs D, Roth HP, Daniel H. Transcriptome and proteome analysis identifies the pathways that increase hepatic lipid accumulation in zinc-deficient rats. J Nutr. 2005;135(2):199–205.

Toporova L, Balaguer P. Nuclear receptors are the major targets of endocrine disrupting chemicals. Mol Cell Endocrinol. 2020;502:110665.

Tovoli F, Stefanini B, Mandrioli D, Mattioli S, Vornoli A, Sgargi D, et al. Exploring occupational toxicant exposures in patients with metabolic dysfunction-associated steatotic liver disease: A prospective pilot study. Dig Liver Dis. 2023 Dec 26:S1590–8658(23)01097-6.

Vilas-Boas V, Gijbels E, Cooreman A, Van Campenhout R, Gustafson E, Leroy K, et al. Industrial, Biocide, and Cosmetic Chemical Inducers of Cholestasis. Chem Res Toxicol. 2019;32(7):1327–1334.

Wahlang B, Appana S, Falkner KC, McClain CJ, Brock G, Cave MC. Insecticide and metal exposures are associated with a surrogate biomarker for non-alcoholic fatty liver disease in the National Health and Nutrition Examination Survey 2003-2004. Environ Sci Pollut Res Int. 2020;27(6):6476–87.

Wang J, Lu P, Xie W. Atypical functions of xenobiotic receptors in lipid and glucose metabolism. Med Rev (Berl). 2022;2(6):611–24.

Yanagisawa R, Koike E, Win-Shwe TT, Yamamoto M, Takano H. Impaired lipid and glucose homeostasis in hexabromocyclododecane-exposed mice fed a high-fat diet. Environ Health Perspect. 2014;122(3):277–83.

Yang C, Zhu L, Kang Q, Lee HK, Li D, Chung ACK, et al. Chronic exposure to tetrabromodiphenyl ether (BDE-47) aggravates hepatic steatosis and liver fibrosis in diet-induced obese mice. J Hazard Mater. 2019;378:120766.

Yang X, Gonzalez FJ, Huang M, Bi H. Nuclear receptors and non-alcoholic fatty liver disease: An update. Liver Research 2020;4(2),88–93.

Zhang D, Wu S, Lan Y, Chen S, Wang Y, Sun Y, et al. Blood manganese and nonalcoholic fatty liver disease: A cohort-based case-control study. Chemosphere. 2022a;287(Pt 4):132316.

Zhang Y, Bao J, Gong X, Shi W, Liu T, Wang X. Transcriptomics and metabolomics revealed the molecular mechanism of the toxic effect of mancozeb on liver of mice. Ecotoxicol Environ Saf. 2022b;243:114003.

Zhao T, Lv WH, Hogstrand C, Zhang DG, Xu YC, Xu YH, et al. Sirt3-Sod2-mROS-Mediated Manganese Triggered Hepatic Mitochondrial Dysfunction and Lipotoxicity in a Freshwater Teleost. Environ Sci Technol. 2022;56(12):8020–33.

Zhao T, Zheng H, Xu JJ, Xu YC, Liu LL, Luo Z. MnO_2_ nanoparticles and MnSO_4_ differentially affected hepatic lipid metabolism through miR-92a/acsl3-dependent de novo lipogenesis in yellow catfish Pelteobagrus fulvidraco. Environ Pollut. 2023;336:122416.

Zuo L, Chen L, Chen X, Liu M, Chen H, Hao G. Pyrethroids exposure induces obesity and cardiometabolic diseases in a sex-different manner. Chemosphere. 2022;291(Pt 2):132935.

