## Supplemental material for "The dithiocarbamate pesticides maneb and mancozeb disturb the metabolism of lipids and xenobiotics in an in vitro model of metabolic dysfunction-associated steatotic liver disease"

**Supplemental Table 1** Information on the 2 donors whose liver pieces were used for the preparation of primary human hepatocytes (PHH)

| **Donor and reference of the PHH batch** | **Sex** | **Age (years)** | **Ethnicity** | **Liver Pathology** | **Other information** |
| --- | --- | --- | --- | --- | --- |
| Donor 1 (PHH 1)  FOI00023050018 | F | 52 | Caucasian | Hepatic mucinous cystadenoma | Not available |
| Donor 2 (PHH 2)  HUM171311A | M | 48 | Caucasian | Colon carcinoma metastasis | HBV, HCV and HIV negative |

**Supplemental Table 2** Primers used in the present study

| **Gene symbol (and alias)** | **Gene name** | **Accession number** | **Forward primer (5’-3’)** | **Reverse primer  (5’-3’)** |
| --- | --- | --- | --- | --- |
| ***AHR*** | Aryl hydrocarbon receptor | NM_001621.5 | GCCGGTGCAGAAAACAGTAA | TAGCCAAACGGTCCAACTCT |
| ***APOB*** | Apolipoprotein B | NM_000384.2 | CCCTCAGTCCTCTCCAGATAAA | GCTGCCTCTTCTTCCCAATTA |
| ***APOC3*** | Apolipoprotein C3 | NM_000040.3 | TGCTCCAGGAACAGAGGT | CCCTGCATGAAGCTGAGAA |
| ***ATF6*** | Activating transcription factor 6 | NM_007348.4 | GTCAGAGAACCAGAGGCTTAAA | CCAACATGCTCATAGGTCCATA |
| ***CD36***  ***(FAT)*** | CD36 molecule  (fatty acid transporter) | NM_001001548.3 | TTGGGAAAGTCACTGCGACA | GGAAATGAGGCTGCATCTGT |
| ***CYP2B6*** | Cytochrome P450 family 2 subfamily B member 6 | NM_000767.5 | CTTCCGGGGATATGGTGTGA | CACTGAGCCTCCTCCTGAAT |
| ***CYP2E1*** | Cytochrome P450 family 2 subfamily E member 1 | NM_000773.4 | TTGAAGCCTCTCGTTGACCC | CGTGGTGGGATACAGCCA |
| ***CYP3A4*** | Cytochrome P450 family 3 subfamily A member 4 | NM_017460.6 | CTTCATCCAATGGACTGCATAAAT | TCCCAAGTATAACACTCTACACACACAA |
| ***DDIT3*  (*CHOP*)** | DNA-damage inducible transcript 3  (C/EBP-Homologous Protein) | NM_001195053.1 | GTCTAAGGCACTGAGCGTATC | CACTTCCTTCTTGAACACTCTCT |
| ***EIF2AK3* (*PERK*)** | Eukaryotic translation initiation factor 2 alpha kinase 3  (Protein kinase R-like ER kinase) | NM_004836.7 | GACCTCAAGCCATCCAACATA | CTGGTCCATTGCAGTCACTAA |
| ***ERN1*  (*IRE1α*)** | Endoplasmic reticulum to nucleus signaling 1  (Inositol-requiring enzyme-1 alpha) | NM_001433.5 | CCAGACAGACCTGCGTAAAT | CCGGTAGTGGTGCTTCTTATT |
| ***HSPA5***  ***(BIP)*** | Heat shock protein family A (Hsp70) member 5  (Binding immunoglobulin protein) | NM_005347.5 | TGGAGGTGGGCAAACAAA | AACTGCATGGGTAACCTTCTT |
| ***MTTP*** | Microsomal triglyceride transfer protein | NM_000253.3 | TACCAGGCTCATCAAGACAAAG | CTGACACCCAAGACCTGATTT |
| ***NR1H4***  ***(FXR)*** | Nuclear receptor subfamily 1 group H member 4  (Farnesoid X-activated receptor) | NM_001206979.2 | TAAAAACGGGGGCAACTGTG | GCCTGTATACATACATTCAGCCA |
| ***NR1I2***  ***(PXR)*** | Nuclear receptor subfamily 1 group I member 2  (Pregnane X receptor) | NM_003889.4 | TCCTTTGCACCGGATTGTTC | TGGACTGCTTGGTGGTAAGT |
| ***NHR1I3***  ***(CAR)*** | Nuclear receptor subfamily 1 group I member 3  (Constitutive androstane receptor) | NM_001077482.3 | CCCCGGGATCGGTTTCT | GTAGGCCTCATTAATGCTCCG |
| ***SLC27A2***  ***(FATP2)*** | Solute carrier family 27 member 2  (Fatty acid transport protein 2) | [NM_003645.4](https://www.ncbi.nlm.nih.gov/entrez/viewer.fcgi?db=nucleotide&id=1519246038) | CCAGAGTCATGGAGGTCTGA | GCTTTTGGAAGACCTGTGGT |
| ***SLC27A4***  ***(FATP4)*** | Solute carrier family 27 member 4  (Fatty acid transport protein 4) | [NM_005094.4](https://www.ncbi.nlm.nih.gov/entrez/viewer.fcgi?db=nucleotide&id=1519499549) | TAAACCCTGCTGAGACCCG | ATTGTGGCCCGGCGT |
| ***TBP*** | TATA-box binding protein | NM_003194.5 | GAGCTGTGATGTGAAGTTTCC | TCTGGGTTTGATCATTCTGTA |
