## Supplemental Figures 1-7 for "The dithiocarbamate pesticides maneb and mancozeb disturb the metabolism of lipids and xenobiotics in an in vitro model of metabolic dysfunction-associated steatotic liver disease"

### ATP levels in HepaRG cells

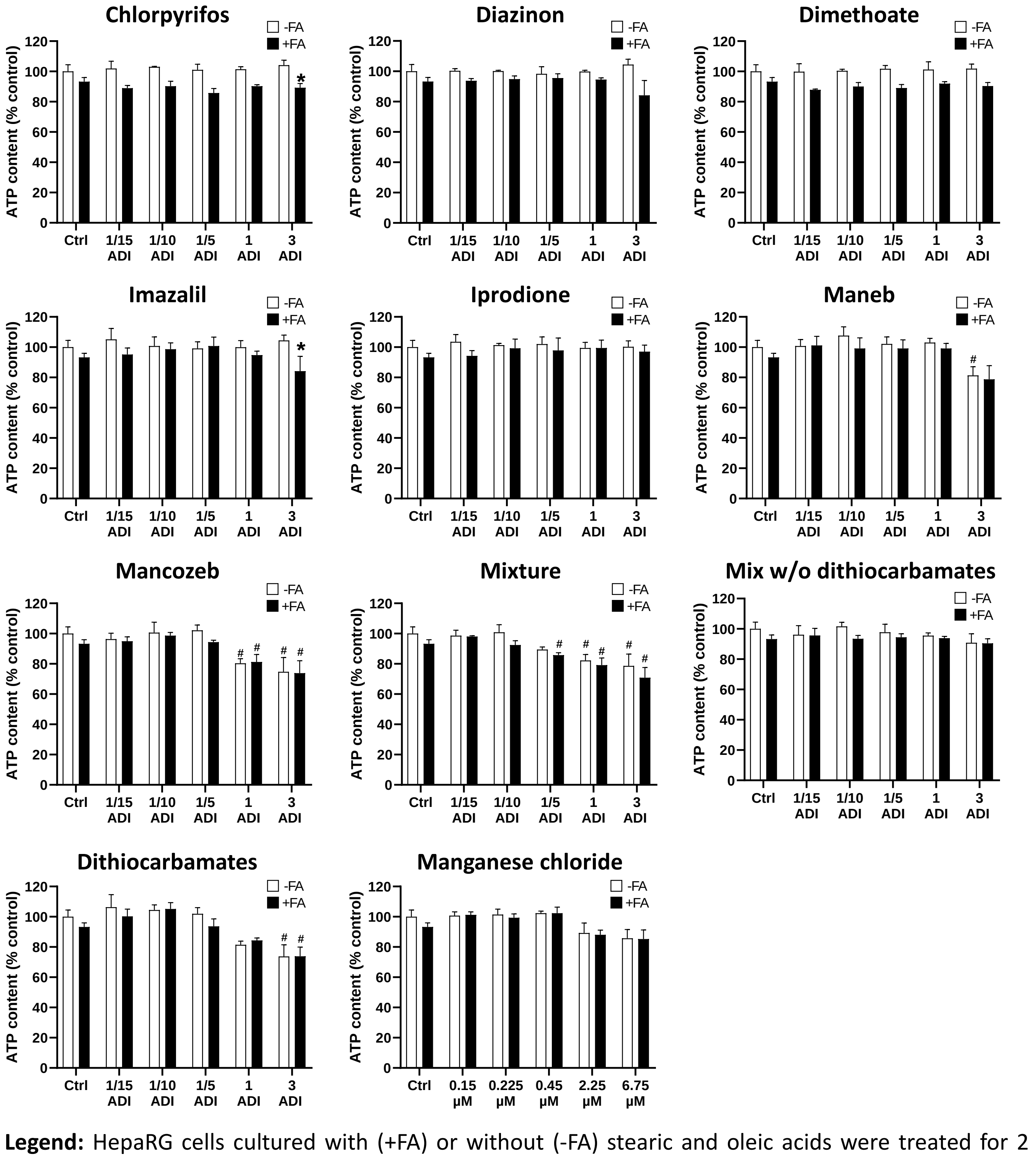


**Legend:** HepaRG cells cultured with (+FA) or without (-FA) stearic and oleic acids were treated for 2 weeks with five different concentrations of chlorpyrifos, diazinon, dimethoate, imazalil, iprodione, maneb, mancozeb, pesticide mixture, pesticide mixture without the dithiocarbamates maneb and mancozeb (mix w/o dithiocarbamates), dithiocarbamates (maneb + mancozeb), or manganese chloride ($MnCl_2$). The different concentrations of $MnCl_2$ (in µM) correspond to the respective ADI concentrations of the dithiocarbamates maneb and mancozeb. ATP levels were assessed after the 2-week treatment as described in the Materials and Methods. Results are means ± SEM for 3 independent cultures and expressed as percentages of the values obtained in untreated -FA-HepaRG cells. *Significantly different from the corresponding treated -FA-HepaRG cells. #Significantly different from the corresponding untreated -FA or +FA-HepaRG cells (Ctrl).

#### Supplemental Figure 1

### Neutral lipids in HepaRG cells

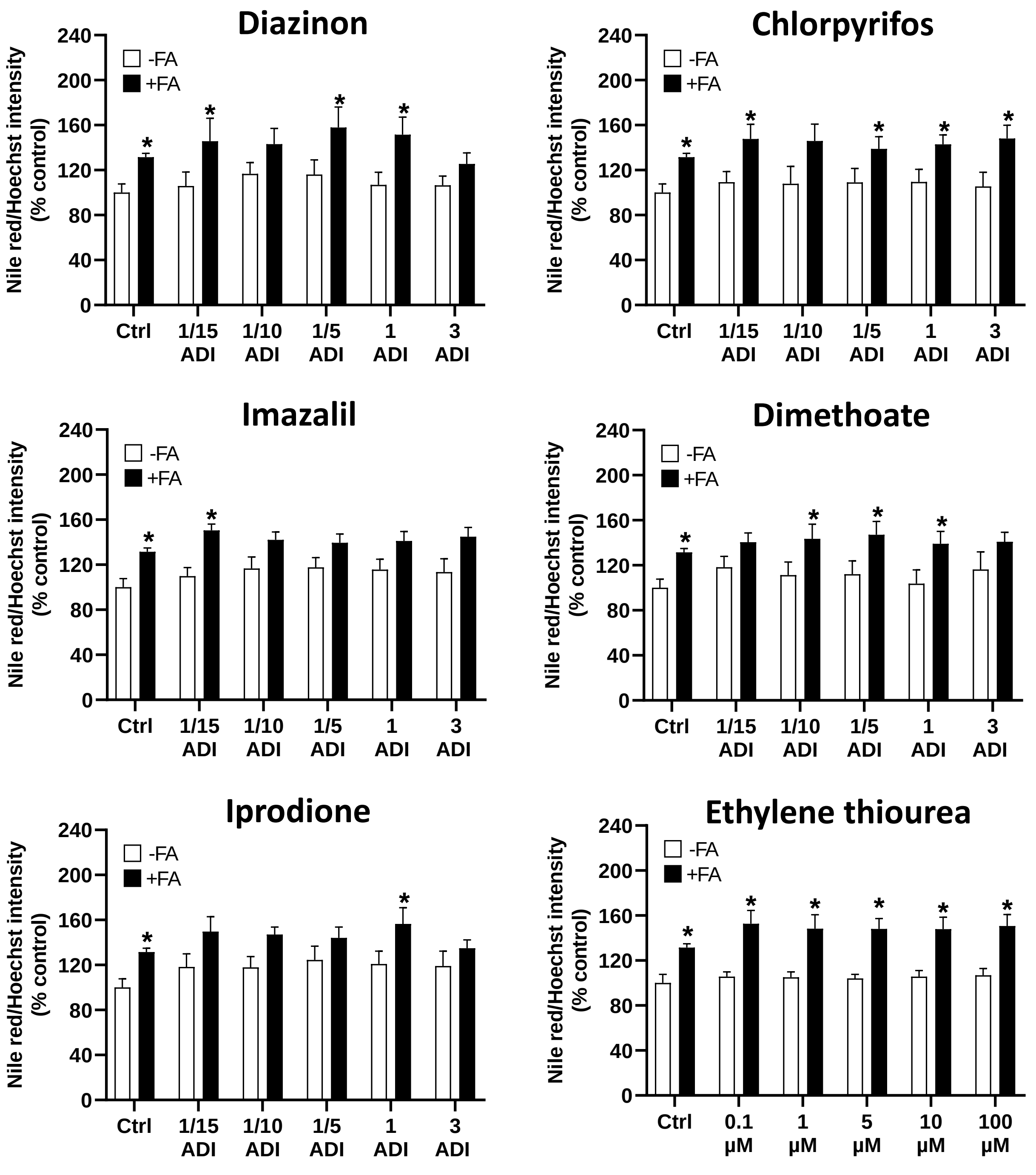


**Legend**: HepaRG cells cultured with (+FA) or without (-FA) stearic and oleic acids were treated for 2 weeks with five different concentrations of diazinon, chlorpyrifos, imazalil, dimethoate, iprodione, or ethylene thiourea, a known metabolite of maneb and mancozeb. The different concentrations of ethylene thiourea (in μM) correspond to the respective ADI concentrations of these dithiocarbamates. After the 2-week treatment, neutral lipids and nuclei were stained with the fluorescent dyes Nile red and Hoechst 33342, respectively and neutral lipids were assessed using the Nile red/Hoechst fluorescence intensity ratio. Results are means ± SEM for 5 to 6 independent cultures and are expressed as percentages of the values obtained in untreated -FA-HepaRG cells. *Significantly different from the corresponding untreated or treated -FA-HepaRG cells.

#### Supplemental Figure 2

### Triglycerides in HepaRG cells

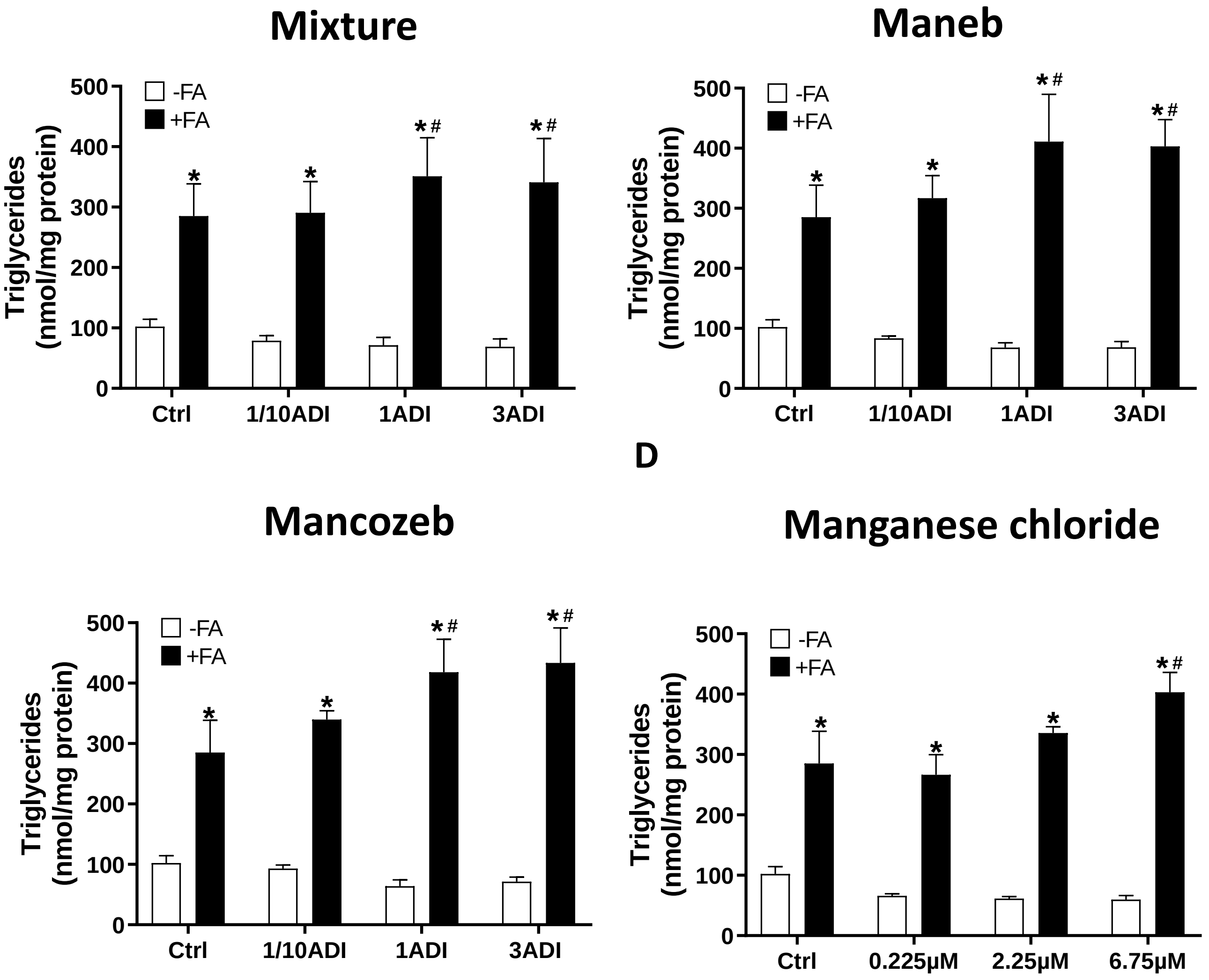


**Legend**: HepaRG cells cultured with (+FA) or without (-FA) stearic and oleic acids were treated for 2 weeks with three different concentrations of the pesticide mixture, maneb, mancozeb, or manganese chloride ($MnCl_2$). The different concentrations of $MnCl_2$ (in $\mu$M) correspond to the respective ADI concentrations of maneb and mancozeb. After the 2-week treatment, cellular triglycerides were assessed as described in the Materials and Methods. Results are means $\pm$ SEM of 3 independent cultures and are expressed in nmol/mg protein. *Significantly different from the corresponding untreated or treated -FA-HepaRG cells. #Significantly different from the corresponding untreated -FA or +FA-HepaRG cells (Ctrl).

#### Supplemental Figure 3

### Mitochondrial fatty acid oxidation in HepaRG cells

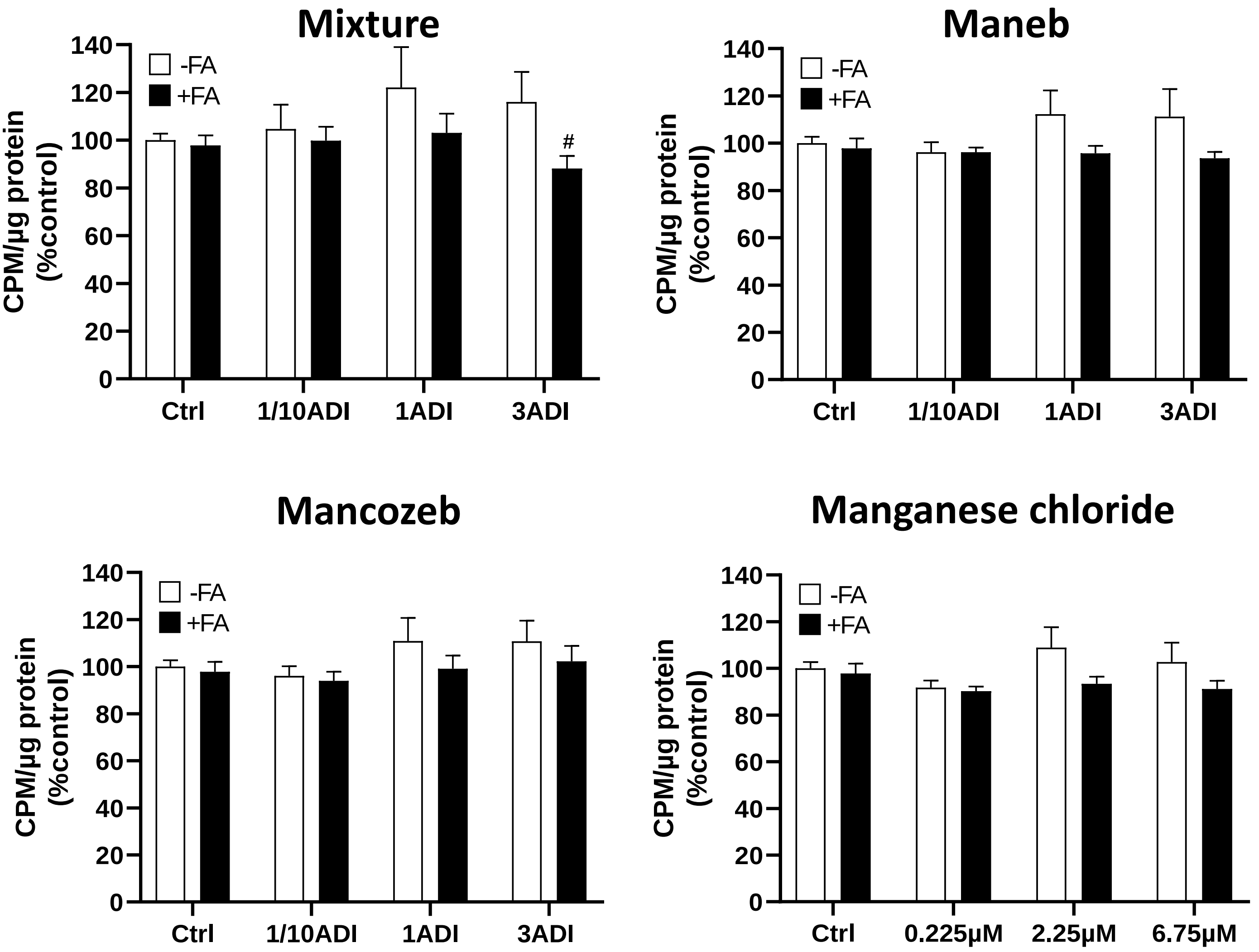


**Legend**: HepaRG cells cultured with (+FA) or without (-FA) stearic and oleic acids were treated for 2 weeks with three different concentrations of the pesticide mixture, maneb, mancozeb, or manganese chloride ($MnCl_2$). The different concentrations of $MnCl_2$ (in $\mu$M) correspond to the respective ADI concentrations of maneb and mancozeb. After the 2-week treatment, mitochondrial fatty acid oxidation was assessed as described in the Materials and Methods. Results are means $\pm$ SEM of 6 independent cultures and expressed as percentages of the values obtained in untreated -FA-HepaRG cells. #Significantly different from the corresponding treated +FA-HepaRG cells (Ctrl).

#### Supplemental Figure 4

### *De novo* lipogenesis in HepaRG cells

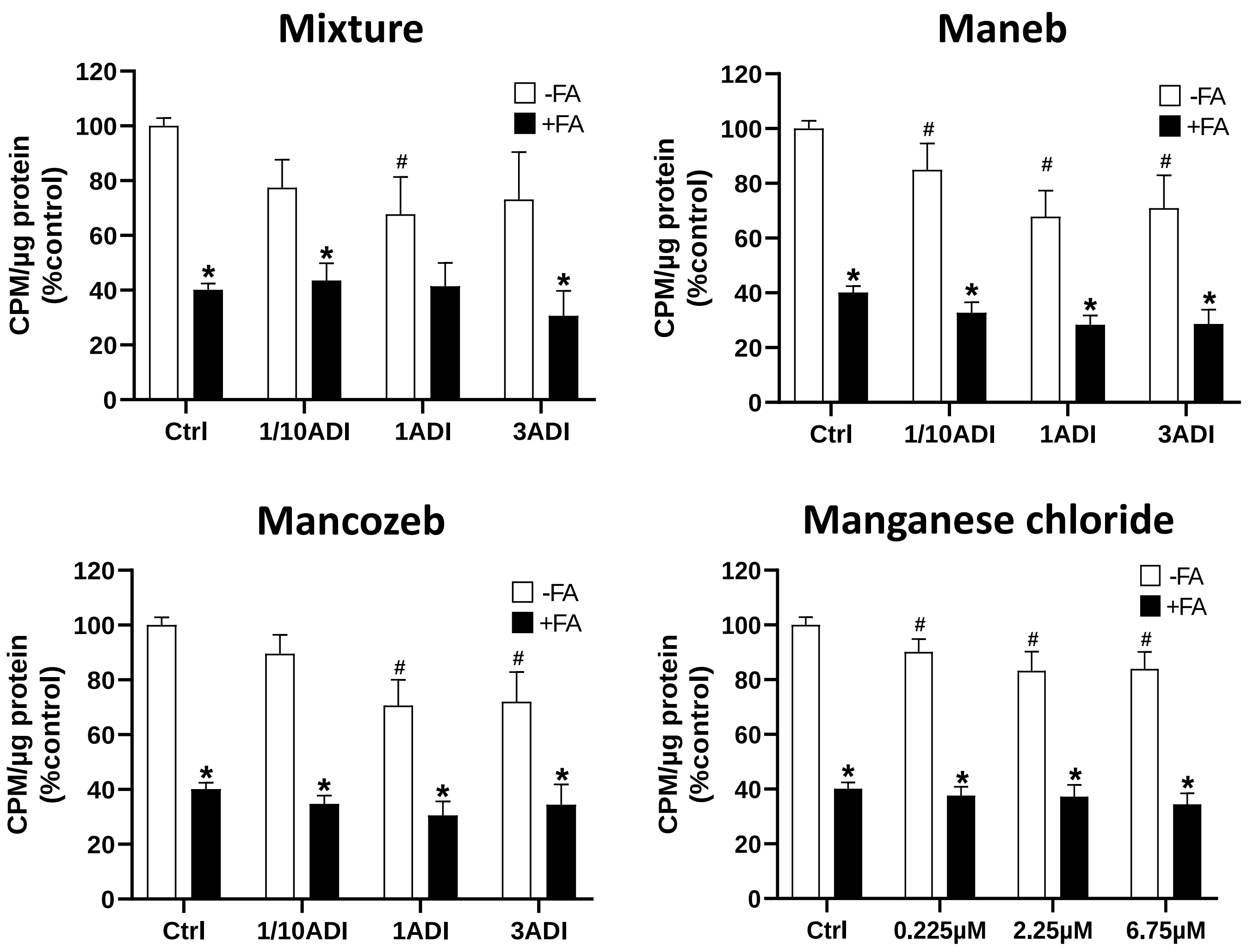


**Legend**: HepaRG cells cultured with (+FA) or without (-FA) stearic and oleic acids were treated for 2 weeks with three different concentrations of the pesticide mixture, maneb, mancozeb, or manganese chloride ($MnCl_2$). The different concentrations of $MnCl_2$ (in $\mu$M) correspond to the respective ADI concentrations of maneb and mancozeb. After the 2-week treatment, *de novo* lipogenesis was assessed as described in the Materials and Methods. Results are means ± SEM of 6 independent cultures and expressed as percentages of the values obtained in untreated -FA-HepaRG cells. *Significantly different from the corresponding untreated or treated -FA-HepaRG cells. #Significantly different from the corresponding untreated -FA-HepaRG cells (Ctrl).

#### Supplemental Figure 5

### Xenobiotic-metabolizing enzyme activity in HepaRG cells

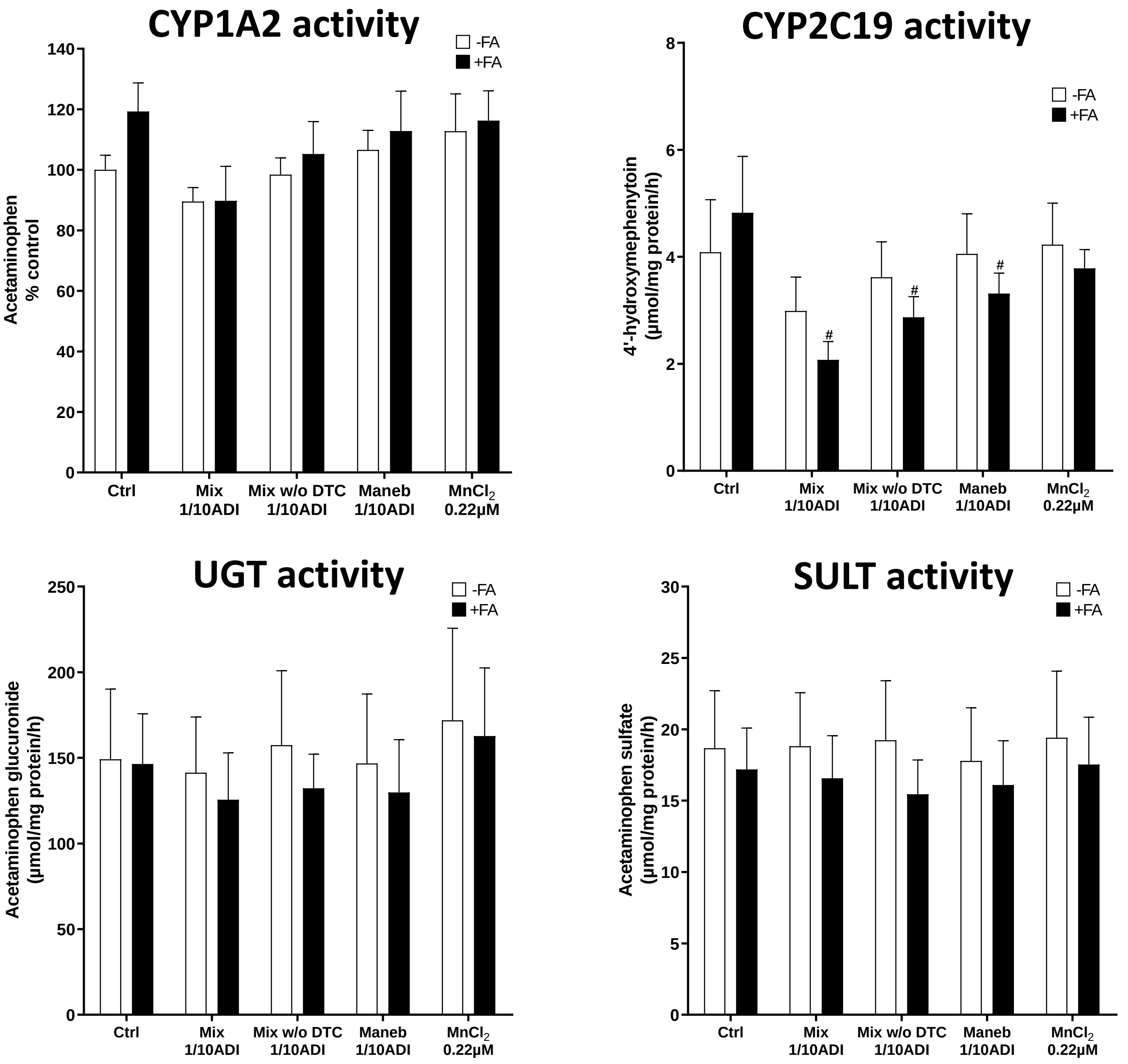


**Legend**: HepaRG cells cultured with (+FA) or without (-FA) stearic and oleic acids were treated for 2 weeks with the pesticide mixture (Mix), the pesticides mixture without the dithiocarbamates maneb and mancozeb (Mix w/o DTC), maneb, or manganese chloride ($MnCl_2$). Cells were treated at the 1/10 ADI concentration for the pesticides, or with 0.22 µM $MnCl_2$, which corresponds to 1/10 ADI of maneb. At the end of the 2-week treatment, cytochrome P450 1A2 (CYP1A2), cytochrome P450 2C19 (CYP2C19), UDP-glucuronosyltransferase (UGT) and sulfotransferase (SULT) activity was measured as described in the Materials and Methods. All enzyme activities are in µmol/mg protein/h (expressed for CYP1A2 activity as percentages of the value obtained in untreated -FA-HepaRG cells). Results are means ± SEM of 5 independent cultures. #Significantly different from the corresponding untreated +FA-HepaRG cells (Ctrl).

#### Supplemental Figure 6

### Chlorpyrifos biotransformation in HepaRG cells

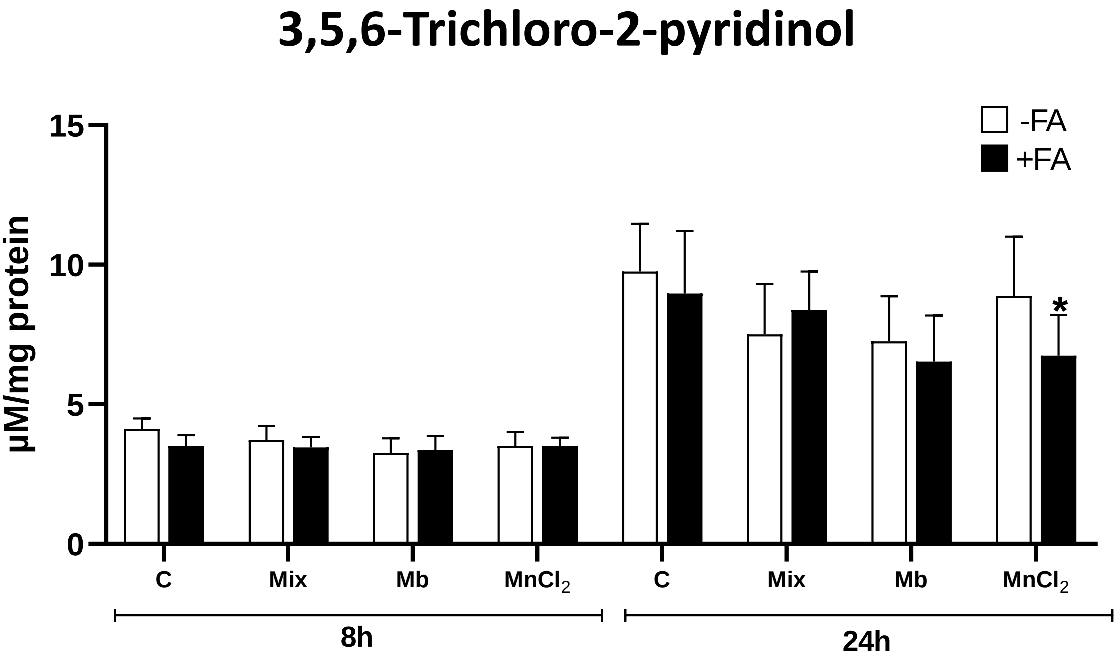


**Legend**: Effects of pesticides and manganese chloride on chlorpyrifos metabolism in differentiated HepaRG cells cultured with or without stearic and oleic acids. HepaRG cells were treated with (+FA) or without (-FA) fatty acids, pesticides, or manganese chloride ($MnCl_2$) for 2 weeks. Cells were treated at the 1/10 ADI concentration for the pesticide mixture (Mix) and maneb (Mb), or with 0.22 µM $MnCl_2$, which corresponds to 1/10 ADI of maneb. After the 2-weeks, HepaRG cells were treated with 50 µM of chlorpyrifos and concentrations of 3,5,6-trichloro-2-pyridinol were measured in the culture media after 8 and 24 h as described in the Materials and Methods. Results are means ± SEM for 4 independent cultures and expressed in µM/mg protein. *Significantly different from the corresponding treated -FA-HepaRG cells.

#### Supplemental Figure 7
